# Active sensory exploration is a major driver of dopamine neuron activity

**DOI:** 10.64898/2025.12.05.692548

**Authors:** Kacper Kondrakiewicz, Xander Temmerman, Sofie Luijten, Len De Paep, Sebastian Haesler

**Affiliations:** VIB-KU Leuven Center for Neuroscience, Leuven, Belgium; Department of Neurosciences, KU Leuven, Leuven, Belgium

## Abstract

Novel stimuli promote arousal, orienting, and learning-related plasticity. Many of these effects have been attributed to dopamine (DA) release, yet whether individual DA neurons encode novelty remains controversial, and the circuit origin of novelty-related signals is unknown. Here, we recorded and manipulated neurons in the ventral tegmental area and substantia nigra (VTA/SNc) of mice exposed to novel and familiar odors while monitoring active sensory exploration, including sniffing and facial movements. Whereas behavior-independent novelty signals were restricted to a small DA subpopulation, most DA neurons tracked sniffing and facial movements, even in the absence of experimentally delivered external stimuli, and irrespective of stimulus novelty. Activity in the mediodorsal pons (mdPons), a major input region to the DAergic midbrain, likewise correlated tightly with sniffing and facial movements and optogenetic stimulation of glutamatergic mdPons neurons induced both sensory exploration and DA firing. Together, these findings identify a circuit mechanism directly linking salient sensory events to DA activity and active sensory exploration.

## Introduction

In many species, novel stimuli elicit arousal, evoke orienting behaviors (Sokolov, 1963; Graham and Clifton, 1966; Modirshanechi et al., 2023) and promote learning (Li et al., 2003). The pervasive influence of novel stimuli on neural function and behavior has been linked to the neuromodulator dopamine (DA). Novelty is associated with DA release in the striatum, and experimentally increasing DA levels can mimic some behavioral and learning effects typically evoked by novelty (Menegas et al., 2017; Morrens et al., 2020; Li et al., 2023; Mocellin et al., 2024; Young et al., 2024; Johnson et al., 2025)

These observations have been incorporated into the influential reward prediction error (RPE) framework of DA function, in which DA activity encodes the discrepancy between expected and received rewards (Schultz et al., 1997). Within this framework, DA neurons treat novel stimuli as a signal of potential reward, giving novel stimuli a ‘novelty bonus’ (Kakade and Dayan, 2002; Gershman and Niv, 2015). Alternatively, DA activation has been proposed to encode other properties of novel stimuli, including their high salience irrespective of value (Horvitz, 2000; Redgrave and Gurney, 2006), the intrinsically rewarding information gain they provide (Bromberg-Martin and Hikosaka, 2009), or the sensory prediction errors they evoke (Gardner et al., 2018).

Although widely accepted, the idea that dopamine (DA) neurons are tuned to novelty has recently been called into question (Monosov, 2024). In non-human primates performing a novelty-seeking task, neither novel stimuli nor cues predicting novelty elicited responses in DA neurons (Ogasawara et al., 2022). Moreover, some studies have reported DA neuron responses to highly familiar sensory stimuli (Horvitz et al., 1997), whereas others found that a subset of DA neurons responded more strongly to familiar than to novel stimuli (Kamiński et al., 2018; see Suppl. Table 1 for a summary of single-neuron findings). Even in studies reporting robust novelty-related activation, such responses are typically observed in only a small fraction of DA neurons (Lak et al., 2016; Kamiński et al., 2018). It remains unclear whether the observed variability across studies reflects differences in experimental protocols or relates to the functional specialization of distinct DA subpopulations. Progress towards resolving this question is complicated by the lack of knowledge about the input projections that drive novelty-related activity in DA cells.

Here, we addressed these questions in head-restrained mice exposed to novel and familiar odors. In rodents, novel odors evoke exploratory sniffing, whisking and orofacial movements (Wesson et al., 2008; Esquivelzeta Rabell et al., 2017; Kurnikova et al., 2017). These orienting reactions largely overlap with active sensing behaviors, through which animals coordinate the sampling of sensory information (Wachowiak, 2011). Hence, we collectively refer to sniffing and orofacial movements as active sensory exploration throughout this study. We measured active sensory exploration both during exposure to novel odors and during spontaneous bouts of active sensing in the absence of any experimentally delivered stimuli. This allowed us to experimentally dissociate novelty-related neural responses from activity associated with sensory exploration. Moreover, each animal underwent classical conditioning to permit the classification of single-units recorded in the VTA/SNc into putative dopaminergic (pDA) and putative GABAergic (pGABA) populations based on their well-established response profiles.

We found that most neurons across both populations were strongly activated by active exploration regardless of stimulus novelty, while only a small subset of pDA neurons exhibited preferential responses to novel stimuli in absence of active exploration. Optogenetic activation and inhibition of DA neurons promoted and suppressed active exploration, respectively. To identify the potential origin of DAergic activity associated with active exploration, we then recorded neurons in two candidate input regions, the mediodorsal pons (mdPons) and superior colliculus. Of these, only the mdPons was tightly coupled to active exploration, and the optogenetic activation of glutamatergic mdPons neurons induced both active exploration and pDA firing. Notably, pDA neurons that responded more strongly to mdPons stimulation were also more strongly modulated by active exploration, yet they were no better at discriminating between novel and familiar stimuli. Together, our findings reveal that a small, distinct pDA subpopulation encoded stimulus novelty independently of behavior, whereas the majority of pDA neurons was driven by active exploration behaviors. This DA subpopulation receives active exploration-related signals through glutamatergic projections from the mdPons, providing a circuit-level explanation for the functional heterogeneity of midbrain DA responses to novel stimuli.

## Results

To study the tuning properties of DA neurons, we performed behavioral experiments in which head-restrained mice sequentially performed i) an odor exposure paradigm and ii) a classical conditioning task. In the odor exposure paradigm, we presented animals with odors in pseudo-random order, while measuring breathing rate, facial movements and pupil diameter. Directly after the odor exposure paradigm, we performed a classical conditioning task, in which two additional odors were paired with a big and a small water reward, respectively (Fig. 1A). During the first 4 experimental days, in the odor exposure paradigm we familiarized animals with a set of 6 odors. On experimental day 5, we introduced 6 novel odors (Fig. 1A), which were interleaved with the familiar odors in pseudo-random order. Novel odors evoked sniffing, facial movements and pupil dilation, as described previously (Wesson et al., 2008; Esquivelzeta Rabell et al., 2017). These components of active exploration habituated over a few trials (Fig. 1B). By experimental day 5, all animals had also rapidly learned the conditioning task. They associated odor cues with the outcome, as indicated by anticipatory licking upon presentation of the odor paired with the big reward (Fig. 1C). Starting from day 5, we repeated a 4-day familiarization period with different odors and introduced novel odors the day after.

**Fig. 1.**
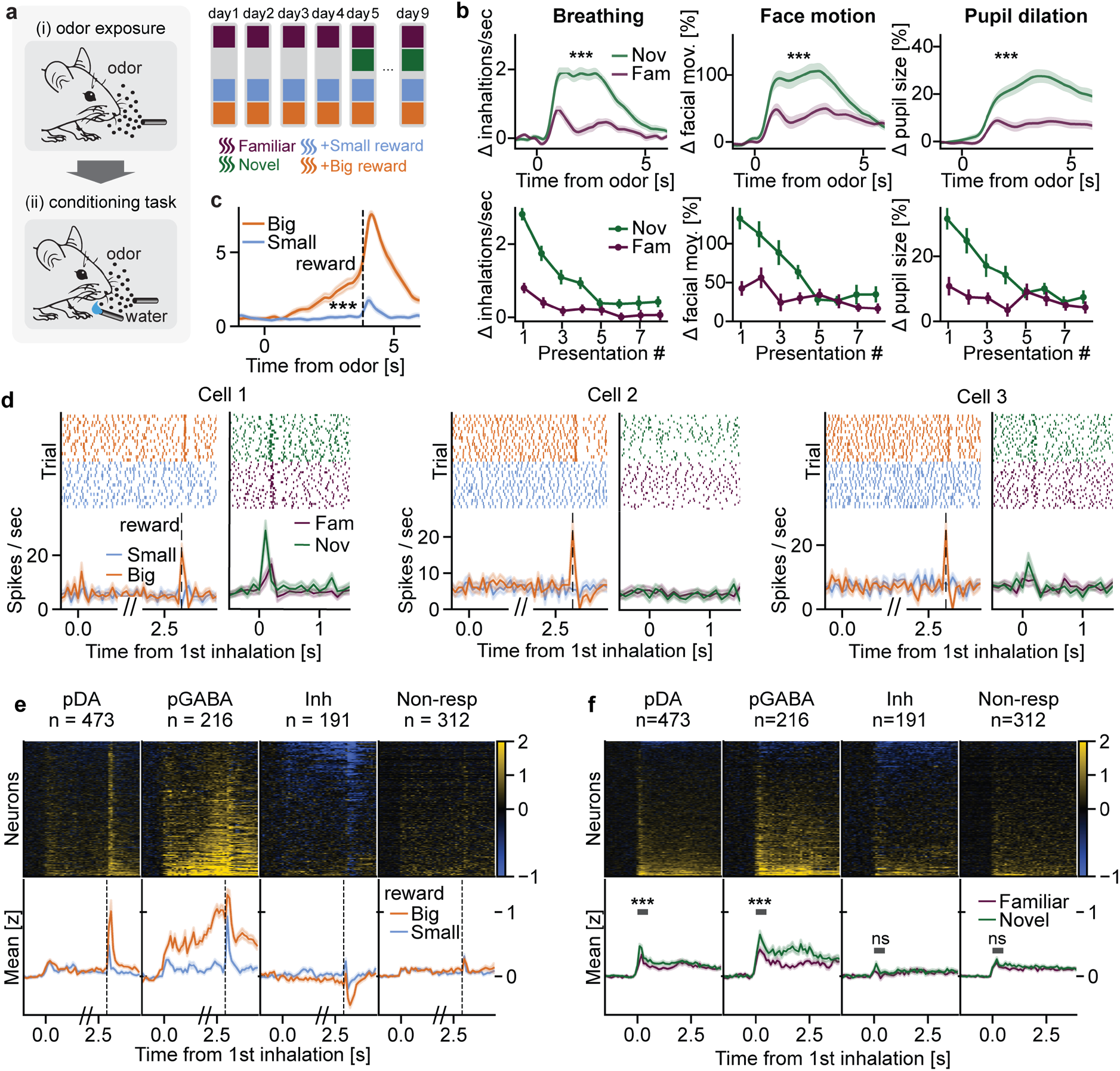
Exposing mice to novel odors triggers active exploration and VTA/SNc activity. **a)** Behavioral protocol. Head-fixed mice (n=26 sessions from 13 animals) were exposed to a set of novel and familiar odorants, while their sniffing rate and facial movements were monitored non-invasively (see Methods). Immediately after, the mice received either big or small water rewards systematically proceeded by previously learnt odor cues. **b)** Behavioral responses to odors. Top: novel odors caused stronger active exploration than familiar ones, as measured by sniffing (t(25)=9.43, p<0.001), facial movements (t(25)=6.45, p<0.001) and pupil dilation (t(25)=5.30, p<0.001; all responses averaged from 1st four presentations). Bottom: habituation curves for novel and familiar odors. **c)** Mice displayed anticipatory licking upon presentation of the big reward cue, which was higher than licking after the small reward cue (t(25)= 5.63, p<0.001). Vertical line: reward delivery. **d)** Responses of three example putative DA neurons. All three cells responded transiently to big rewards, but responses to novel stimuli were much more variable. **e)** Responses to cues and rewards from all recorded cells, split into putative types. Top: z-scored responses of individual cells in big reward trials. Traces – population average in big reward (orange) and small reward (blue) trials. **f)** Responses of the same neurons (sorted separately) to novel and familiar stimuli (heatmap for novel). Only pDA and pGABA populations responded stronger to novel than to familiar odorants (W=36467.5, p<0.001 and W=6950.5, p<0.001, respectively; average from 1^st^ four presentations in 0-500ms window).

On the days when novel odors were presented, we recorded extracellular activity from the ventral tegmental area (VTA, 1008 single-units), substantia nigra pars compacta (SNc, 105 single-units) and retrorubal field (RRF, 79 single units), using chronically implanted 4-shank Neuropixels 2.0 probes (Suppl. Fig. 1, van Daal et al., 2021). It was previously shown that neurons in VTA/SNc have distinct response properties to conditioned stimuli (CS) and rewards (unconditioned stimulus, US), which can be used to reliably identify DA and GABA neurons (Cohen et al., 2012). We used this approach to cluster all recorded cells into putative cell types (Fig 1D, Suppl. Fig. 1B). As expected, we found cells which were transiently activated by CS and/or US (putative DA, pDA; see examples on Fig. 1D), cells with ramping activity between CS and US (putative GABA, pGABA; examples on Suppl. Fig. 1C) and cells showing a sustained decrease in firing between CS and US (*Inh*.; Fig. 1E). A fraction of recorded neurons did not show any responses in the conditioning task (*Non-resp*).

In the odor exposure paradigm, only pDA and pGABA populations demonstrated robust activation by odors (Fig. 1F, Suppl. Fig. 1D). The remaining types (*Inh* and *Non-resp*) did not show significant odor responses. While on average, pDA and pGABA neurons responded more strongly to novel than to familiar odors, we observed considerable variability in the population activity. Less than a third of pDA and pGABA cells responded to novel odors (18% and 28%, respectively; Fig. 1D, 1F), and even the responsive ones were often only activated in a subset of trials (Fig. 2A).

**Fig. 2.**
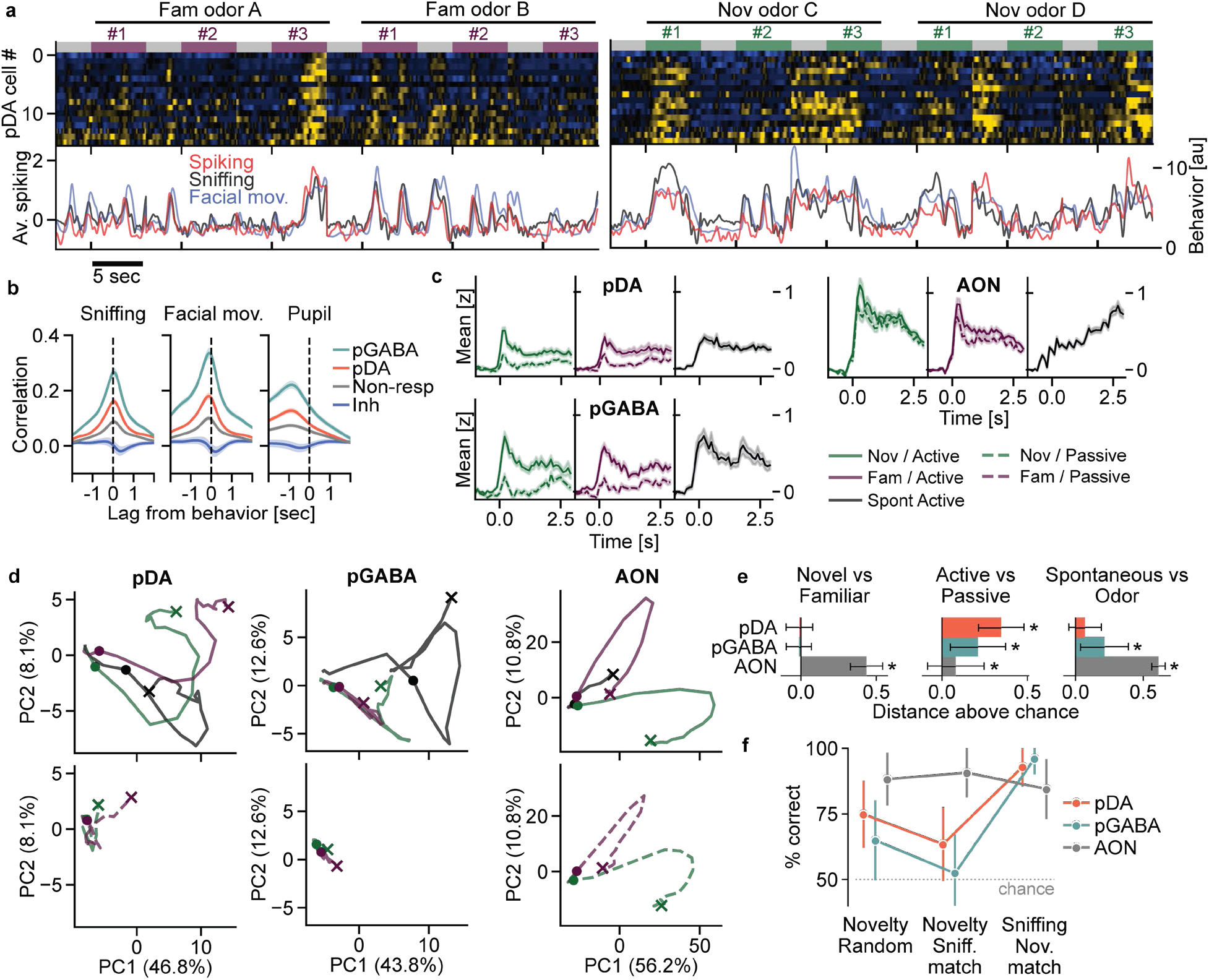
Active exploration dominates pDA and pGABA population activity. **a)** Example activity of all pDA neurons recorded during a single session (n=16). The firing rate (average in red, lower panel) tracked sniffing rate (black) and face motion (blue). Trials from the 1^st^ three presentations of the same odorants were concatenated for illustration purposes. The horizontal bars on top indicate different time epochs: baseline before odor presentation (gray), familiar odors delivery (purple) and novel odors delivery (green). **b)** Activity of pDA and pGABA populations was correlated in time with all three measures of active exploration: sniffing, face motion and pupil dilation. **c)** Mean z-scored responses of pDA (n = 473 neurons, 26 sessions), pGABA (n = 216 neurons, 25 sessions) and AON neurons (n = 1053 neurons, 11 sessions) to novel (green) and familiar (purple) odors, accompanied by active exploration (solid lines) or not (dashed lines; mean ± SEM across neurons; for single-neuron heatmaps, see Suppl. Fig. 2E). Responses (mean z-score 0–500 ms after the first inhalation) were analyzed separately for each population with linear mixed models (fixed effects: novelty, sniffing; random intercepts: session and neuron; likelihood-ratio tests). In pDA neurons, responses were higher in active than passive trials (χ²(1) = 169.9, p < 0.001), and the effect of novelty depended on sniffing (novelty × sniffing: χ²(1) = 4.7, p = 0.030): novel odors evoked stronger responses than familiar odors during active (p < 0.001) but not passive trials (p = 0.57). pGABA responses were higher during active trials (χ²(1) = 200.8, p < 0.001) and unaffected by novelty (novelty: χ²(1) = 0.3, p = 0.57; novelty × sniffing: χ²(1) = 0.8, p = 0.37). AON responses were higher for novel than familiar odors (χ²(1) = 20.8, p < 0.001), both during active (p = 0.002) and passive trials (p = 0.011). They were also higher during active than passive trials (χ²(1) = 55.8, p < 0.001), with no novelty × sniffing interaction (χ²(1) = 1.6, p = 0.20). Responses to spontaneous bouts of exploration (solid black lines) were plotted for comparison, but not included in the statistical model. Post hoc p-values are FDR-corrected. **d)** Population trajectories in pDA and pGABA, but not AON, were determined mainly by active exploration. Note the similar shape of AON trajectories between active vs. passive conditions. Active – solid lines, Passive – dashed lines; Novel – green, Familiar – purple, Spontaneous – black. Dot – odor delivery or exploration onset (in the Spontaneous condition), cross – trajectory end. **e)** pDA and pGABA population activity distinguished between Active vs. Passive, but not Familiar vs. Novel conditions. The normalized distances were obtained by subtracting average distance between random subsets of trials. The standard errors were obtained by recalculating distances using data from a random half of mice (see Methods). **f)** Decoding novelty from pDA/pGABA activity decreased when controlling for the sniffing level (pDA: t(299) = −10.46, p < 0.001; pGABA: t(299) = −10.47, p < 0.001). In contrast, in AON, controlling for sniffing led to slightly better decoding of stimulus novelty (t(299) = 2.95, p = 0.003). Decoding the sniffing level (below vs. above average rate, novelty-matched) was more accurate than decoding novelty (sniffing-matched) in pDA (t(299) = 32.37, p < 0.001) and pGABA (t(299) = 46.75, p < 0.001), whereas in AON the opposite effect was found (t(299) = −7.16, p < 0.001). Points: mean ± SD across 300 resampling iterations. Dotted line: chance level.

Fig. 2A shows the activity of all the pDA neurons recorded simultaneously in one mouse during a single session in response to 2 familiar and 2 novel stimuli, overlaid with the sniffing rate and facial movement. As illustrated, pDA cells increased activity predominantly during active exploration, which sometimes occurred spontaneously (without odors). To quantify this relationship, we calculated the cross-correlations between the measured behaviors and neuronal activity (Fig. 2B). The firing rate of pDA and pGABA cells was on average positively correlated with both sniffing and facial movements, and only to a lesser degree with pupil dilation (which was also much slower, lagging behind neurons ∼0.8 sec). pDA and pGABA activity was correlated with respiration rate only during epochs of active exploration, i.e. during sniffing (>3Hz) but not during epochs of slower baseline breathing (<3Hz, Suppl. Fig. 2A-B).

To investigate how active exploration and stimulus novelty jointly contributed to pDA and pGABA population activity, we split all odor-presentation trials into four categories: novel vs. familiar × active exploration vs. passive stimulus reception (Fig. 2C). Because the behavioral measures of active exploration were highly redundant (Suppl. Fig. 2C), we classified trials as active vs. passive based solely on the presence or absence of a sniffing response (see Methods). Using the same criteria, we also detected episodes of active exploration that occurred spontaneously during baseline periods before odor presentation. To make conditions comparable, trials included in the analysis were selected to have equivalent sniffing levels (e.g., between novel and familiar sets; see Suppl. Fig. 2D).

We first computed the mean response of each neuron across conditions (Fig. 2C). In both pDA and pGABA neurons, responses were dominated by exploration: neuronal activity was much higher in active than in passive trials (Fig. 2C). Novelty had a smaller influence. pDA neurons responded slightly more strongly to novel than to familiar odors, but only when these were actively explored, whereas pGABA responses did not depend on novelty in either condition (Fig. 2C).

To control for the possibility that these results reflected the experimental design rather than intrinsic pDA/pGABA response properties, we repeated the same analysis on a previously published dataset containing responses from the olfactory cortex (anterior olfactory nucleus, AON) recorded using the same odor exposure paradigm (de Plus et al., 2025). In contrast to pDA and pGABA neurons, AON neurons responded robustly also during trials not accompanied by active exploration, and novel odors evoked stronger responses than familiar odors regardless of whether they were actively explored (Fig. 2C, right).

To examine these effects at the population level, we reduced the dimensionality of the population responses using principal component analysis (PCA). The resulting state-space trajectories for each condition are illustrated in Fig. 2D. Responses of both pDA and pGABA populations differentiated robustly between active exploration and passive reception of sensory stimuli (Fig. 2D–E). In contrast, the difference in population activity between novel and familiar stimuli did not exceed baseline, estimated by shuffling trial labels between conditions (Fig. 2D–E; see Methods). Additionally, pDA, but not pGABA, activity was relatively similar during spontaneous and odor-evoked active exploration. In contrast, AON population activity robustly separated novel from familiar stimuli. The separation between active and passive trials was smaller, but also statistically significant (Fig. 2D–E, right).

Taken together, these results indicated that the activity of both pDA and pGABA populations was dominated by responses to active exploration. To further test this conclusion, we trained an SVM classifier on population activity. The activity of the pDA population predicted whether a stimulus was novel or familiar with 75% accuracy, but only when a random subset of trials was used (Fig. 2F). When the trials used for decoding were selected to have similar sniffing rates in the novel and familiar conditions (*sniffing-matched*), the performance dropped significantly, to 63%. In contrast, a classifier trained to distinguish between trials with high vs. low sniffing reached the highest accuracy (93%), even when the proportion of novel and familiar stimuli in each condition was matched (*novelty-matched*). For the pGABA population, we observed qualitatively similar results: novelty decoding dropped from 65% to 52% when sniffing was matched, whereas sniffing was decoded with 96% accuracy. In contrast, decoding from AON was stable across conditions: stimulus novelty could be decoded with ∼90% accuracy both from random and from sniffing-matched trial sets, whereas decoding sniffing was slightly less accurate (84%; Fig. 2F).

Although pDA and pGABA population activity was dominated by active exploration, the above analyses did not exclude the possibility that individual neurons encoded novelty. We therefore next examined the relationship between different factors regulating neuronal activity at the single-cell level. The degree of correlation with active exploration varied substantially across single cells, both during odor presentations and during baseline periods (Fig. 3A). Typically, the same neurons were consistently modulated by all three aspects of active exploration, namely sniffing, facial movements, and pupil dilation (Fig. 3B, Suppl. Fig. 3A), as expected given that these behaviors usually occurred together (Suppl. Fig. 2C). More importantly, neurons that were more strongly modulated by active exploration exhibited also stronger responses upon odor presentations in general (Fig. 3B). However, the modulation by active exploration was independent of how strongly a cell differentiated between novel and familiar stimuli. Consistent with this result, we observed two types of responses among individual pDA cells: some neurons were regulated primarily by active exploration, whereas others were regulated primarily by stimulus novelty (see examples in Fig. 3C).

**Fig. 3.**
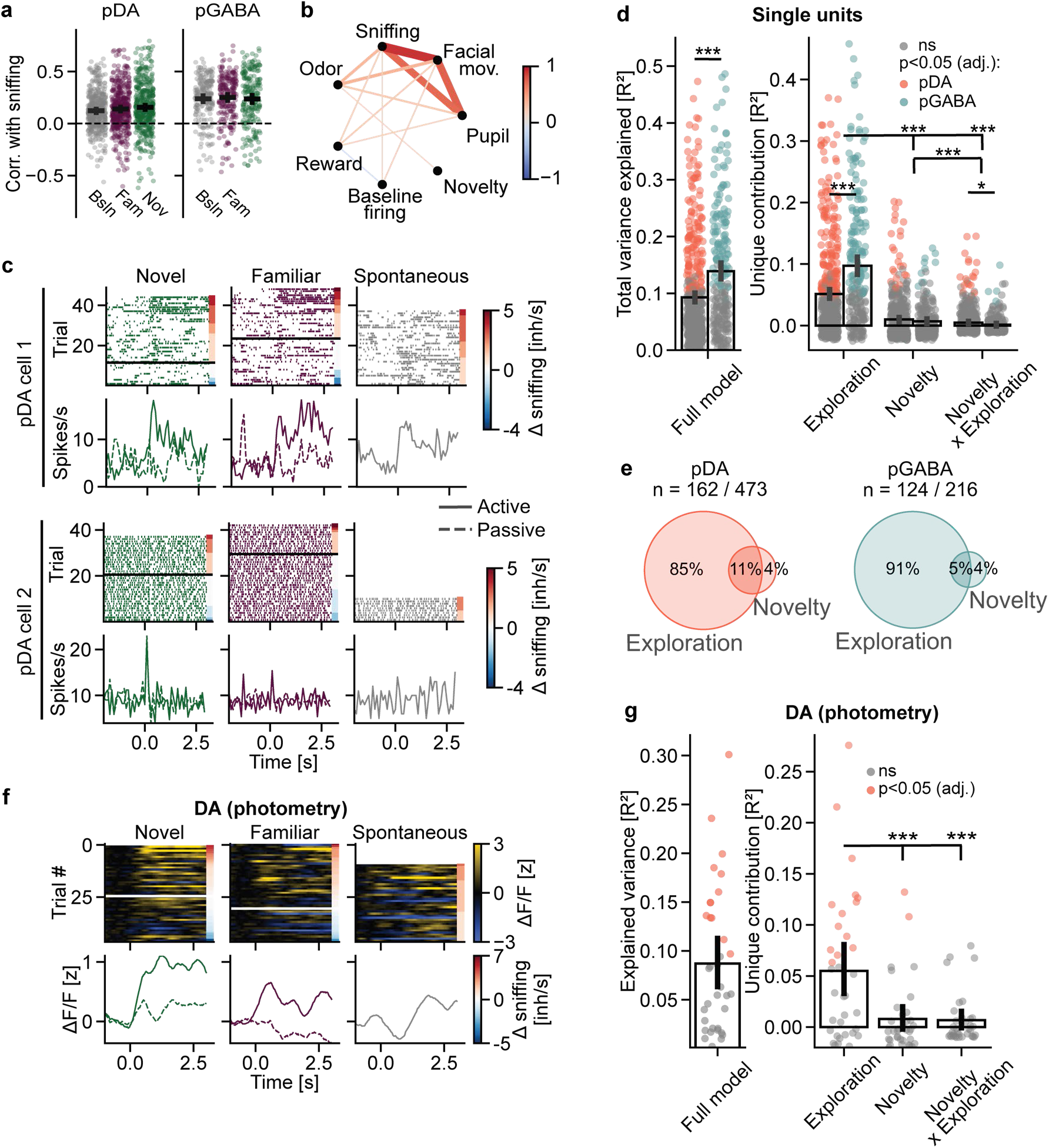
Most DA single-units are activated by active exploration. **a)** The distribution of correlations with sniffing for all recorded pDA and pGABA cells. The correlations (0 lag) were computed separately for 3 epochs: baseline (gray), familiar odors presentations (purple) and novel odors presentations (green). **b)** Correlation structure between different response properties of individual pDA cells. The correlations with sniffing, facial movements and pupil dilation were computed using data from baseline (pre-odor) periods. Cells more strongly correlated with sniffing were also more strongly correlated with facial movements (r_s_=0.90, p<0.001) and pupil size (r_s_=0.72, p<0.001); correlations with facial movements and pupil size were similarly related (r_s_=0.75, p<0.001). Cells more strongly modulated by active exploration had higher responses to odors in general (sniffing: r_s_=0.32; facial movements: r_s_=0.35; pupil: r_s_=0.22; all p<0.001), but did not differentiate more strongly between novel and familiar odors (sniffing: r_s_=−0.07, p=0.13; facial movements: r_s_=−0.10, p=0.07; pupil: r_s_=−0.08, p=0.15). Correlation strength is represented by both line width and color (red for positive, blue for negative). Only correlations with p<0.05 (FDR adjusted for multiple comparisons) are shown. **c)** Examples of two representative pDA cells – one regulated primarily by active exploration (top), and another one primarily by novelty (bottom). Top panels: raster plots illustrating responses in individual trials. The colored rectangles on the right indicate the magnitude of sniffing response observed in each trial (compared to baseline before odor presentation). Horizontal black line: the split between Active vs. Passive conditions. Bottom panels: average response in each trial category (Active – solid line, Passive – dashed line; Novel – green, Familiar – purple, Spontaneous – black). **d)** Prediction of pDA and pGABA responses with linear regression based on sniffing and novelty. Left: general prediction level (R^2^), right: contribution of individual predictors. Single-trial pDA and pGABA activity was predicted better by the magnitude of sniffing responses than by stimulus novelty (F(2,1374)= 322.13, p<0.001; post hoc: t(688)=16.44, p<0.001) or the novelty × sniffing interaction (t(688)=18.82, p<0.001). The contribution of sniffing was larger in pGABA than in pDA cells (cell type × predictor interaction: F(2,1374)=47.48, p<0.001; post hoc: t(324)=5.82, p<0.001). Each predictor’s unique contribution is the mean decrease in R² after shuffling it across trials (100 permutations). Dots: single cells; colored dots: shuffling the predictor significantly reduced prediction accuracy for that cell (permutation test, p<0.05, FDR-corrected), bars: mean from all cells, error bars: standard errors of the mean. **e)** A small fraction of pDA and pGABA cells were regulated exclusively by odor novelty (rather than sniffing). **f)** Example of DA calcium traces recorded from a single fiber. Top panels: heatmaps illustrating responses in individual trials. The colored rectangles on the right indicate the magnitude of sniffing response observed in each trial (compared to baseline before odor presentation). Horizontal white line: the split between Active vs. Passive conditions. Bottom panels: average response in each trial category (Active – solid line, Passive – dashed line; Novel – green, Familiar – purple, Spontaneous – black). **g)** Single-trial calcium responses of the DA population were predicted better by the sniffing rate than by stimulus novelty (F(2,68)=16.72, p<0.001; post hoc: t(34)=4.01, p<0.001) or the novelty × sniffing interaction (t(34)=4.30, p<0.001). Individual dots represent single fibers (n=35 fibers from 8 mice); colored dots: shuffling the predictor significantly reduced prediction accuracy for that fiber (permutation test, p<0.05, FDR-corrected). Insert: example calcium trace (red) overlaid with sniffing rate (black).

To characterize this diversity quantitatively, we fitted a regression model predicting average firing rate of each cell in every trial based on active exploration (measured with sniffing), stimulus novelty and interaction between the two (Fig. 3D). Shuffling individual predictors revealed that both pDA and pGABA cells were regulated predominantly by active exploration. The impact of stimulus novelty was overall smaller (Fig. 3D), but still significant for a fraction pDA neurons. Among the pDA neurons for which the regression model achieved significant prediction (162 out of 473), 85% of cells were regulated exclusively by exploration, 4% exclusively by novelty and 11% by both (Fig. 3E). Within the pGABA population, similar proportions were found (91%, 4% and 5%, respectively).

To ensure that our conclusions were not biased by errors in cell type classification, we repeated our analysis on calcium transients recorded with fiber photometry from genetically identified DA neurons (Fig. 3F, Supl. Fig. 4A-B). The results were strikingly similar and indicated that DA neurons were modulated predominantly by exploration rather than by stimulus novelty (Fig. 3G). Moreover, we ruled out the possibility that the relationship between sniffing and neuronal activity could be related to differences between individual odorants (Suppl. Fig. 4C). We also we confirmed there was no stronger modulation by novelty in the short time window just after odor delivery (<100msec), when active exploration responses had not yet been initiated (Suppl. Fig. 4D-E). Finally, we tested other measures of active exploration (facial movements and pupil dilation) and found similar results as when using sniffing (Suppl. Fig. 4G).

**Fig. 4.**
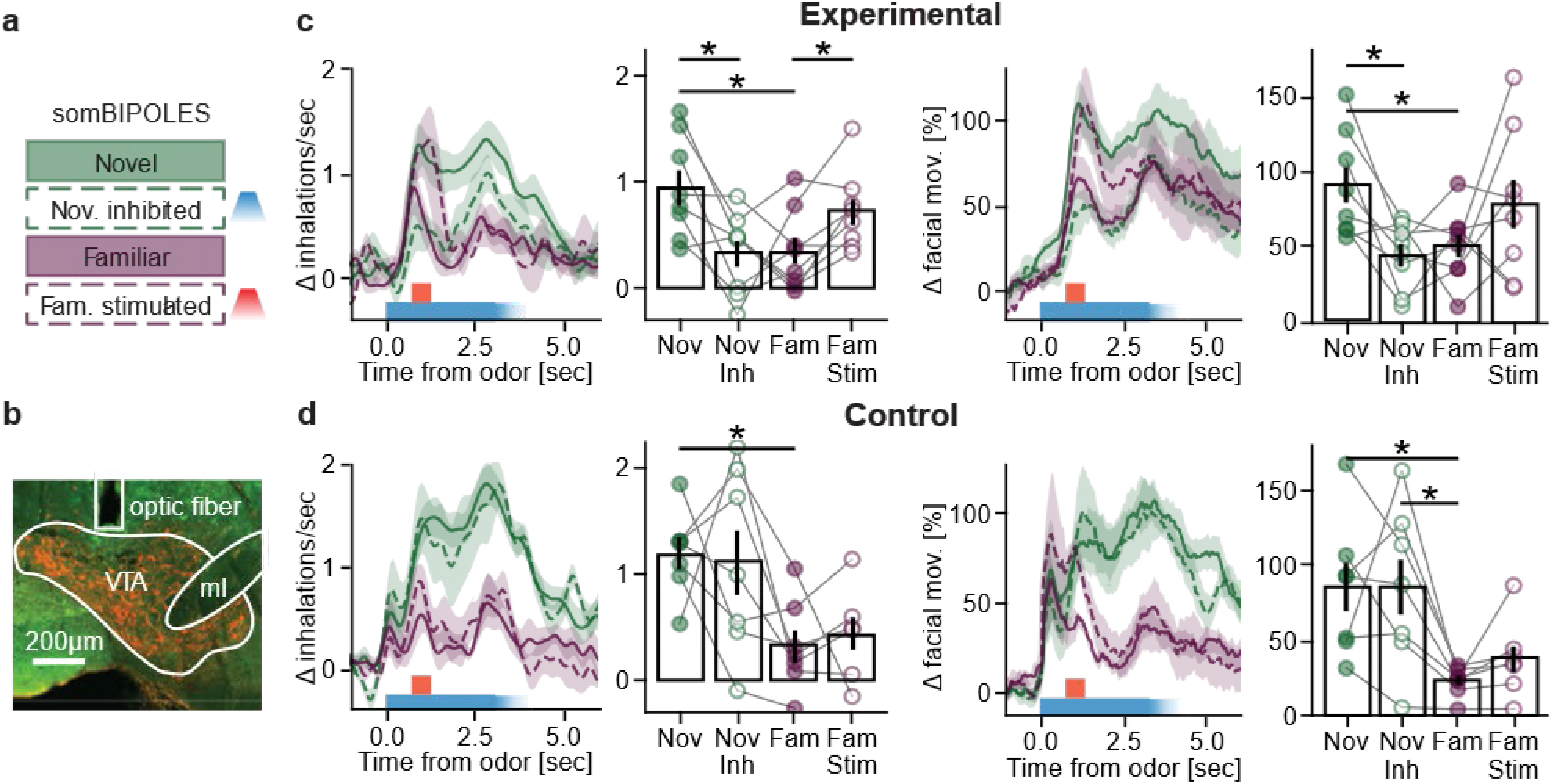
Stimulation of VTA dopamine neurons promotes active exploration, inhibition suppresses it. **a)** Experimental design. A subset of novel odors (green) was paired with blue-light inhibition (dashed lines). A subset of familiar odors (purple) was paired with red-light stimulation (dashed lines). **b)** Example expression of optogenetic construct (AAV5-hSyn-DIO-somBIPOLES-mCerulean, red) in the VTA of DAT-Cre mouse. **c)** In the experimental group (DAT-Cre, n=8) novel odors evoked more sniffing (F(3,21)=4.96, p<0.01; post-hoc: t(8)=3.41, p<0.05) and facial movements (F(3,21)=3.13, p<0.05, post-hoc: t(8)=3.83, p<0.05) than familiar odors. Optogenetic inhibition of DA neurons reduced novelty-evoked orienting (sniffing: t(8)=3.35, p<0.05, facial movements: t(8)=3.44, p<0.05), while optogenetic stimulation increased sniffing to familiar stimuli (t(8)=3.33, p<0.05, facial movements: t(8)=1.37, ns.). **d)** In the control group (wild-type, n=7) novel odors evoked more sniffing (F(3,15)=4.26, p<0.05; post-hoc: t(7)=4.70, p<0.05) and facial movements (F(3,15)=4.26, p<0.01, post-hoc: t(7)=4.05, p<0.05) than familiar odors, but optogenetic stimulation did not cause any behavioral changes (all p>0.1).

Next, we tested whether the activity of DA neurons was causally related to active exploration. We focused on DA cells in VTA, where most of our recorded cells originated from (Suppl. Fig. 1). Optogenetic inhibition of this population (Fig. 4A-B) during presentation of novel odors reduced the sniffing responses and facial movements to the level typically observed for familiar stimuli (Fig 4C). Brief activation of DA neurons during presentation of familiar odors increased sniffing rate (Fig. 4C). The effects on pupil dilation followed a similar trend but were not statistically significant (Suppl. Fig. 5). No effects of the optogenetic stimulation could be observed in the control group (Fig 4D). Taken together, these results indicate that DA activity is necessary and sufficient to evoke at least some components of the active exploration typically caused by stimulus novelty.

Since the degree of modulation by active exploration varied greatly among pDA neurons, we asked whether these differences might be explained by distinct input patterns to individual DA cells. Specifically, we hypothesized that a collection of nuclei in mediodorsal pons (mdPons), including the lateral parabrachial nucleus, the laterodorsal tegmental nucleus and locus coeruleus, might convey signals related to active exploration (for more anatomical information, see Suppl. Fig 6). All these nuclei send monosynaptic projections to both DA and GABA midbrain neurons (Watabe-Uchida et al., 2012) and collectively regulate arousal in response to salient sensory stimuli (Fuller et al., 2011). Specifically, mdPons areas project to premotor nuclei that control different aspects of active explortaion: the intermediate reticular zone (sniffing and facial movements, (Deschênes et al., 2012; Yang et al., 2020; Biancardi et al., 2023)) and Edinger Westphal nucleus (pupil dilation, (Cornwall et al., 1990; Wang and Munoz, 2015)). As an alternative candidate input region, we studied the superior colliculus (SC), which also targets VTA/SNc and is implicated in active exploration (Allen et al., 2021; Solié et al., 2022a).

First, we recorded electrophysiological activity in both regions from mice exposed to novel/familiar odors (mdPons: 810 single-units, SC: 172 single-units, Fig. 5A). Cells from mdPons displayed stronger correlation with sniffing, facial movements and pupil dilation than the ones from SC (Fig. 5B & Suppl. Fig. 6). In the odor exposure paradigm, population responses in both regions were modulated stronger by active exploration than by novelty (Fig. 5C). However, the effect of active exploration was stronger in mdPons than in SC (Fig. 5C). Correspondingly, regression analysis indicated that most modulated cells in both regions were regulated by exploration, with smaller fraction of cells affected by novelty (Fig. 5D). The overall prediction of single-trial activity was better in mdPons than in SC (Fig. 5D). Finally, neurons positively modulated by exploration were not clustered in a particular nucleus within mdPons, but were spread across the whole recorded area (Fig. 5E; see Suppl. Fig. 6B for the results split into individual structures). Taken together, the results suggested that mdPons was a better candidate for activating DA system in the context of active exploration than SC.

**Fig. 5.**
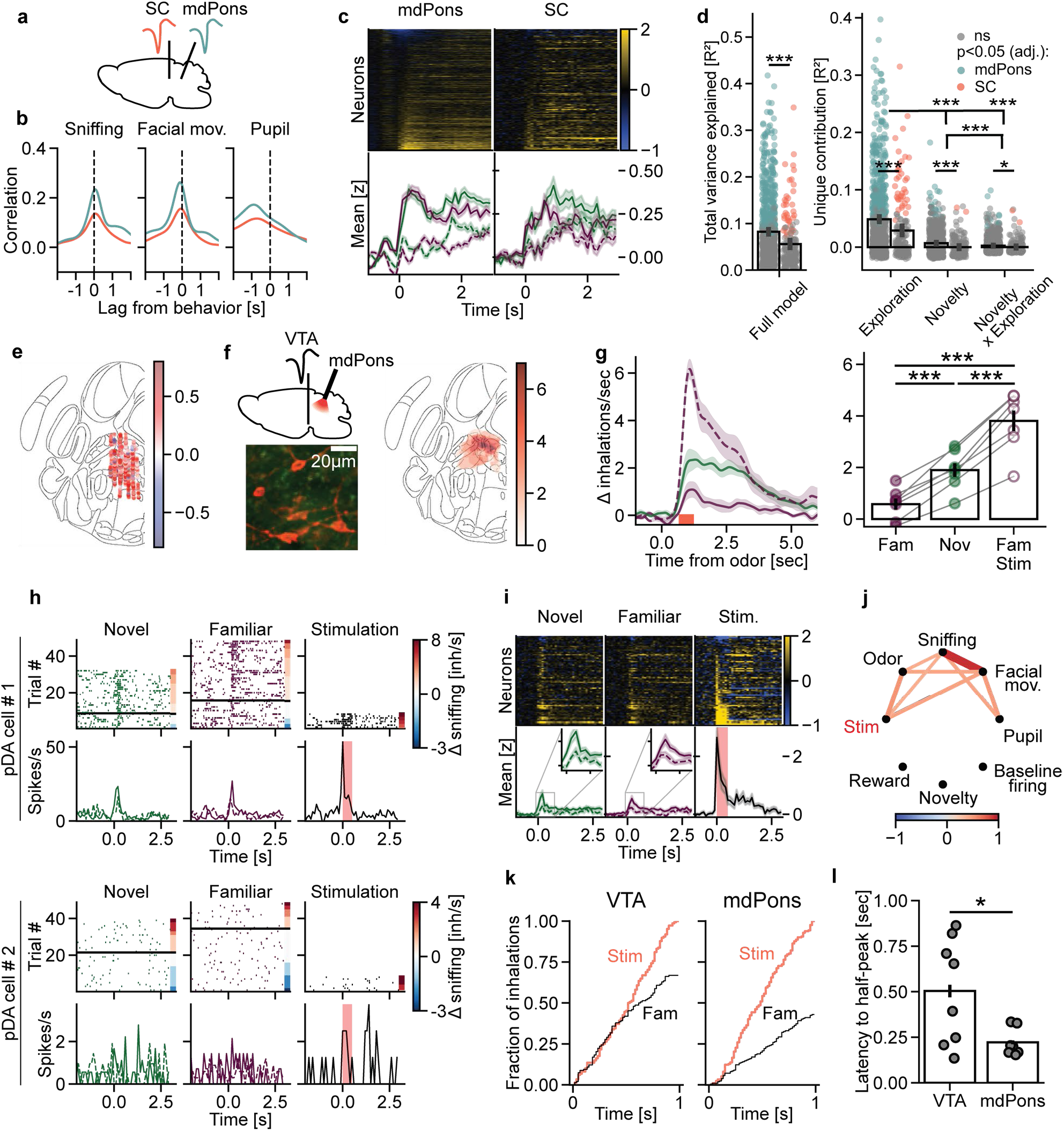
Glutamatergic projections from mediodorsal pons drive both active exploration and pDA activity. **a)** Single-unit recordings from superior colliculus (SC) or mediodorsal pons (mdPons) performed in awake mice (n=12 sessions from 6 mice). **b)** The average cross-correlations between firing rate and orienting-related behaviors, plotted separately for mdPons (green) and SC (red). **c)** Population responses were more strongly modulated by active exploration than by odor novelty in both mdPons (n=810 cells from 4 mice) and SC (n=172 from 2 mice). In mdPons, sniffing exerted a strong effect (χ²(1) = 422.42, p < 0.0001), exceeding the effect of novelty (χ²(1) = 1.29, p = 0.256), with a significant novelty × sniffing interaction (χ²(1) = 18.60, p < 0.0001). In SC, a significant main effect of sniffing was also observed (LMM, χ²(1) = 59.34, p < 0.0001), whereas neither novelty (χ²(1) = 1.09, p = 0.297) nor the novelty × sniffing interaction (χ²(1) = 0.68, p = 1.0) were significant. A combined model confirmed that the influence of active exploration differed between regions (population × sniffing interaction: χ²(1) = 21.26, p < 0.0001), indicating stronger exploration-related modulation in mdPons than in SC. **d)** Single-trial responses of mdPons could be predicted better than responses of SC (U = 85978, p < 0.001). Neuronal activity was explained better by sniffing than by odor novelty or by the novelty × sniffing interaction (repeated-measures ANOVA: F(2,1960) = 373.30, p < 0.001; post hoc: Sniffing vs. Novelty, t(981) = 18.21, p < 0.001; Sniffing vs. Novelty × Sniffing, t(981) = 20.92, p < 0.001; Novelty × Sniffing vs. Novelty, t(981) = 6.55, p < 0.001). Post hoc comparisons further revealed that both sniffing (t(300) = 4.12, p < 0.001) and novelty (t(303) = 4.66, p < 0.001) had bigger influence of single-neuron responses in mdPons than in SC. **e)** Spatial distribution of the recorded mdPons cells vs. their correlation with sniffing (positive correlations: red, negative: blue). For illustration purposes, cells recorded from different antero-posterior locations (-5.6 to -5.0mm from bregma) are plotted on a single atlas plane. **f)** Top left: experimental design. Recording of VTA electrophysiological activity accompanied by the optogenetic stimulation of mdPons (n=7). Bottom left: example expression of the optogenetic construct (AAV5-hSyn-DIO-somBiPOLES-mCerulean, red) in mdPons of VGlut2-Cre mice. Right: expression of the construct summarized for all injected animals. The darkest color indicates overlapping expression in all 7 animals. **g)** Average of behavioral effects (n=7). Brief optogenetic stimulation of glutamatergic mdPons cells evoked higher sniffing response than familiar (F(2,12)=101.12, p<0.001; post hoc: t(6)=10.94, p<0.001) or novel (t(6)=9.01, p<0.001) odors. **h)** Example responses of two pDA cells recorded during the mdPons stimulation experiment. Only cells which were strongly activated by the mdPons stimulation (top) displayed excitatory responses to novel/familiar odors (bottom). Green: novel odors, purple: familiar odors, black: no odor. Dashed lines: mean from passive trials. **i)** Average responses of all pDA cells recorded in the stimulation experiment (n = 78 from 7 mice), after presentation of novel odors (green), familiar odors (purple), or mdPons stimulation (black, no odor). Heatmaps on top illustrate responses across neurons in the active exploration trials (for novel/familiar odors) or across all trials (for the optogenetic stimulation). **j)** Correlation structure between response properties of pDA cells recorded in the stimulation experiment. Cells which were more strongly activated by the optogenetic stimulation were also more correlated with sniffing during baseline (pre-odor) periods (r_s_=0.39, p<0.01) and had higher responses to odors in general (r_s_=0.53, p<0.001), but did not differentiate stronger between novel vs. familiar odors (r_s_=0.08, p=0.69) nor had higher baseline firing rate (r_s_=0.12, p=0.48). Only correlations with p<0.05 (FDR adjusted for multiple comparisons) are shown. **k)** Sniffing dynamics in response to the optogenetic stimulation of VTA or mdPons, plotted for two representative animals. The plot illustrates cumulative distribution of inhalations across time after the presentation of familiar odorant (aligned to first inhalation), quantified separately for the control condition (no stimulation, in black) and experimental condition (optogenetic stimulation, in red). **l)** The latency of sniffing responses to optogenetic stimulation, quantified with the ZETA test (see Methods) for each animal. The latency was longer for mdPons VGlut2 stimulation than for VTA DA stimulation, t(8.09)=2.63, p<0.05.

To test if the input form mdPons could evoke both active exploration and DA firing, we optogenetically stimulated glutamatergic cells from this brain region, while recording VTA/SNc activity as described above (Fig. 5F). Brief stimulation of mdPons during presentation of familiar odors triggered active exploration that were even stronger than typically evoked by novel stimuli, including sniffing (Fig 5G), facial movements and pupil dilation (Suppl. Fig. 7). No effects were found in the control group (Supl. Fig. 7). The stimulation of mdPons caused also a strong activation of pDA neurons (Fig. 5H-I), consistently with previous reports (Lodge and Grace, 2006). The pDA cells with the strongest responses to optogenetic stimulation were also more modulated by active exploration (also before odor delivery) and showed higher responses upon presentation of odors (Fig. 5J; see examples on Fig. 5H). These results could not be accounted for by general excitability, because the responsiveness to mdPons stimulation was not significantly related to how strongly pDA neurons were activated by rewards, nor to their baseline firing rate (Fig. 5J). Importantly, the responsiveness to mdPons stimulation was also not related to how strongly pDA cells differentiated between novel vs. familiar odors. We concluded that the subpopulation of pDA neurons activated by active exploration received particularly strong excitatory input from mdPons. The results of mdPons inhibition were more variable (Suppl. Fig. 8) and in some animals caused paradoxically an increase rather than decrease in exploration. However, the effects on pDA neurons were in each case consistent with the directionality of the behavioral effect (that is, exploration and pDA firing either decreased or increased together).

Since optogenetic stimulation of both VGlut neurons in the mdPons and DA neurons in the VTA neurons evoked active exploration, we asked if the two structures were part of the same pathway, or if they belonged to two distinct pathways for evoking behavioral responses. VTA DA neurons have no direct projections to respiratory control centers but can influence sniffing rate through a polysynaptic pathway involving the ventral striatum and corticothalamic loops (Li et al., 2023; Johnson et al., 2025). mdPons, on the other hand, could trigger sniffing indirectly through the VTA, but also directly through projections to the reticular zone (Yang et al., 2020; Biancardi et al., 2023; Kaur et al., 2024), a collection of premotor nuclei that control sniffing and facial movements (Deschênes et al., 2012). When we compared the behavioral response latencies after optogenetic stimulation, we found that mdPons stimulation evoked exploration much faster than DA stimulation (Fig. 5K-L), consistently with a model predicting recruitment of a direct pathway, independent of the VTA.

## Discussion

In this study, we examined how the sensation of novelty is linked to active exploration and to the activity of DA and non-DA neurons in the VTA/SNc. We exposed head-fixed mice to novel and familiar odors while monitoring sniffing, facial movements and pupil diameter. Although pDA and pGABA neurons responded on average more strongly to novel than to familiar odors, this difference was largely accounted for by the exploratory behaviors that novel odors evoked. Only a small fraction of pDA neurons encoded novelty *per se*, irrespective of exploratory behavior. The majority of pDA neurons tracked exploratory behaviors on a trial-by-trial basis, regardless of whether they were evoked by novel odors or occurred spontaneously, and novelty could be decoded less accurately once exploratory behaviors were matched across conditions. Bidirectional optogenetic manipulations showed that VTA DA neurons not only tracked exploration but also contributed to generating it. Finally, we identified the mdPons as an input region whose activity was tightly coupled to exploration, and whose glutamatergic neurons, when activated, evoked both exploratory behavior and pDA firing.

A key implication of our findings is that they help reconcile conflicting observations about the encoding of novelty in DA neurons. While several studies reported DA responses to novel stimuli (Steinfels et al., 1983; Ljungberg et al., 1992; Kamiński et al., 2018), a recent report found no such responses when the stimuli were not associated with reward (Ogasawara et al., 2022). Our novelty paradigm did not include any external rewards, yet it revealed a small fraction of novelty-responsive pDA neurons. There might thus indeed be a small DA subpopulation encoding stimulus novelty independently of other task variables, but whose identification may require extensive sampling. On the other hand, the vast majority of pDA neurons was associated with active exploration, supporting a role for dopamine in the approach and exploration of novel stimuli or states, as had been suggested earlier (Kakade and Dayan, 2002).

Novel, surprising or social stimuli evoke stronger sniffing and orofacial movements than familiar, expected or non-social stimuli (Wesson et al., 2008; Solié et al., 2022b). Unless these and other spontaneous, uninstructed behaviors are carefully monitored and accounted for, neural responses can easily be misattributed to stimulus properties. Accordingly, some previously reported responses to novel (Lak et al., 2016) or otherwise salient stimuli (Horvitz, 2000; Comoli et al., 2003) may therefore, at least in part, reflect active exploratory behaviors. This interpretation might also account for seemingly paradoxical observations, such as DA responses to familiar visual stimuli (Kamiński et al., 2018), DA activation during approach of familiar social partners (Dai et al., 2022; Solié et al., 2022a), and the activation of punishment-responsive DA neurons by novel stimuli, which is difficult to reconcile with a uniform novelty bonus signal (Kakade and Dayan, 2002; Akiti et al., 2022). However, whether a wider repertoire of exploratory behaviors in freely moving animals drives DA activity in a similar way as observed in this study remains to be tested.

To identify the origin of exploration-related activity in the VTA/SNc, we examined two major input regions, the SC and mdPons. Both were modulated by active exploration, but this modulation was considerably stronger in the mdPons. The weaker coupling in the SC may reflect its role in spatial aspects of orienting, such as saccades and head and body turns (Pradel et al., 2021; Solié et al., 2022a), which were constrained in our head-fixed paradigm. Optogenetic activation of glutamatergic mdPons neurons evoked both active exploration and pDA firing (Fig. 5G–I), consistent with earlier reports of pontine excitation of DA neurons (Lodge and Grace, 2006). Moreover, pDA neurons most strongly activated by mdPons stimulation were also those most strongly modulated by active exploration and most responsive to odors in general (Fig. 5H–J). However, they did not discriminate better between novel and familiar odors. Thus, the strength of mdPons input can explain much of the heterogeneity in exploration-related DA activity, but not the novelty selectivity of the small subpopulation described above.

While our optogenetic experiments demonstrated that both the stimulation of glutamatergic neurons of the mdPons and the stimulation of DA neurons evokes active exploration, the analysis of optogenetic response latencies uncovered critical differences between the two structures. Activation of glutamatergic mdPons neurons resulted in short-latency facial movements and sniffing, which is consistent with their anatomical projection pattern (Fig. 5K-L, Suppl. Fig. 9). Excitatory mdPons neurons directly project to the intermediate reticular zone (Yang et al., 2020; Biancardi et al., 2023) – a brain region controlling both breathing and facial movements (Deschênes et al., 2012) – as well as to the Edinger Westphal nucleus which controls pupil dilation (Cornwall et al., 1990; Wang and Munoz, 2015). On the other hand, the response latencies after DA activation were comparatively longer (Fig. 5K-L), consistent with an indirect motor activation through a ventral striatal pathway described recently (Li et al., 2023; Johnson et al., 2025). These insights support a model in which the mdPons serves as an orienting hub, that drives orienting and dopamine release through its excitatory projections (Suppl. Fig. 9). Further research is needed to identify the input pathway to the mdPons and characterize the specific output channels by which different mdPons nuclei control active exploration and DA firing, respectively.

A possible limitation of our study is that individual cells were classified as putative DA or GABA based on their response properties in a classical condition task described previously (Cohen et al., 2012). Therefore, some neurons might have been misclassified. However, we confirmed the main results by selectively measuring activity in DA neurons with fiber photometry (Fig. 3F). Moreover, even potential misclassifications would not change our general conclusions, since vast majority of both pDA and pGABA cells were regulated by active exploration rather than by stimulus novelty (Fig. 3E).

Another possible confound could be that both DA neuron activity and active exploration were driven by the appetitive value of individual odorants. However, our data strongly speaks against this interpretation. The behavior-related modulation of DA activity was observed even in trials with the same odorant (Fig. 2A) and controlling for odorant identity in the regression model did not diminish the effects of exploration (Suppl. Fig. 4). Moreover, active exploration and dopaminergic activity were tightly correlated even in the absence of externally provided sensory stimuli (Fig. 2C), consistently with previous observations (Johnson et al., 2025). While many DA neurons were thus primarily modulated by spontaneous or stimulus-evoked active exploration, we cannot rule out that they additionally encoded value-related or sensory information (Kahnt and Schoenbaum, 2025).

Finally, an alternative interpretation of the limited DA activity during trials not accompanied by active exploration is that the animals failed to detect odor novelty. We consider this explanation highly unlikely, given that the olfactory cortex neurons recorded previously in the same task (de Plus et al., 2025) were robustly modulated by stimulus novelty irrespective of whether novel stimuli were followed by active exploration or not (Fig. 2C-F; see also de Plus et al., 2025; Schiltz et al., 2026).

Taken together, our results indicate that in the context of novelty, the activity of most VTA/SNc DA neurons reflects active exploration rather than novelty itself, and that this exploration-related drive is conveyed, at least partially, by glutamatergic projections from the mdPons. Novelty *per se* is encoded by a small DA subpopulation, whose inputs remain to be identified. Through exploration-related activation, novel and other salient stimuli can evoke DA release in target regions associated with learning (Lisman and Grace, 2005; Morrens et al., 2020) without requiring individual DA neurons to be tuned to novelty. This mechanism fits a broader framework of dopaminergic regulation of motor and autonomic function (Engelhard et al., 2019; Markowitz et al., 2023; Yin, 2023), and helps explain how salient sensory stimuli drive physiological reactions, exploration and memory formation.

## Methods

### Animals

All procedures were executed according to the guidelines set forth by the Federation of European Laboratory Animal Science Associations (FELASA) and received approval from the Animal Ethics Committee at KU Leuven (protocol 161/2021). In total, 69 male mice of the following strains were used: DAT-IRES-Cre (Jackson identifier: #006660, n = 42), VGlut2-IRES-Cre (#016963, n=12), ChAT-IRES-Cre (#006410, n=9), C57Bl/6j (#000664, n=6). All animals were obtained from KU Leuven animal facility, weighed between 20 and 40g and were between 2 and 6 months old.

### Surgical procedures

#### a) General procedure

All experimental animals were surgically implanted with a titanium head plate to permit head restraining during behavioral experiments. Mice were anesthetized with an intraperitoneal injection of a mixture of ketamine (75mg/kg) and medetomidine (Domitor, 1mg/kg) and fixated on a feedback-controlled heating pad at 38°C in a stereotactic set-up (SR-6M-HT, Narishige). 50µl of Lidocaine (Xylocain) was injected subdermally for local anesthesia. A patch of skin on top of the skull was removed, and the edges of the exposed tissue were covered with surgical glue (Vetbond). A dental drill (Athena Champion AC 5000) with a carbide drill head of 0.8mm diameter (FG1, NeoBurr) was used to roughen the skull. Finally, a head plate was fixated on the skull with a 3-component dental cement (Super-Bond, Sun Medical). The anesthesia was reversed using Atipamezole (Antisedan, 0.5mg/kg, IP injection), immediately followed by a dose of Meloxicam as analgesic (Metacam, 5-10 mg/kg, IP).

#### b) Neuropixels 2.0 implantation

The Neuropixels 2.0 probes were prepared, chronically implanted and recovered (1-4 times) as described previously (van Daal et al., 2021). Briefly, after glueing probes to protective cases, the shanks were coated with a fluorescent dye (DiI) and surrounded with a wall built from a mixture of bone wax and mineral oil (50:50). After exposing the skull as described above, a ground screw was inserted above the left visual cortex (contralateral to the probe). The head plate implantation was performed as described above, but the part of skull above the target structure was left exposed. Craniotomy was performed in this place using the following coordinates: for VTA - AP +1.0mm measured from lambda, ML 0. 25 to1.0; for SC - AP 0.0 to 0.75 from lambda, ML 0.9; for brainstem - AP –1.5 from lambda, ML: 0.25 to 1.0 (in the latter case the probe was implanted with 15 degrees angle to the front). Next, duratomy was performed and the probe was inserted (4.75mm deep for VTA, 3.5mm for the two other targets) using a motorized micromanipulator (World Precision Instruments, cat.# DC3314R), with a typical insertion speed of 10µm/s. The probe was fixed to the skull using additional layer of the dental cement (Super Bond, Sun Medical).

#### c) Fiber photometry - surgery

For the fiber photometry experiments, viral vector (AAV9-CAG-FLEX-jGCaMP8s-WPRE, Addgene) was injected unilaterally into VTA/SNc of DAT-Cre animals using a glass capillary (800nL split into 4 injections: ML: 0.5 & 1.2mm, V: 4.5 & 4.0 from brain surface, AP: 1.0 from lambda), with a maximum speed of ∼100nL/min. After the injection, an array of 12 optic fibers (Neurophotometrics) was implanted into VTA/SNc. Individual fibers were 5.0/5.3 mm long (for odd and even fibers, respectively) and 100µm thick, with inter-fiber spacing of 0.25mm (for production protocol, see (Sych et al., 2019)). The array was implanted at the following coordinates: AP 1.0mm from lambda, V 4.2 (for longer fibers) from the brain surface, ML from ∼0. 2 to ∼2.2. The rest of the surgery was performed as described in the section ‘General procedure’.

#### d) Optogenetics – surgery

For the optogenetic manipulations, AAV5-hSyn-DIO-somBiPOLES-mCerulean vector was injected (for VTA: 350 nL/hemisphere, AP 1.1mm from lambda, ML 0.7, V 4.5; for brainstem: 200-250 nL/hemisphere, AP: -1.5 from lambda with 15 degrees angle to the front, ML 1.2 with 15 degrees angle to the middle, V 2.5). After that, optic fiber cannulas (#33835, Netiona; fiber FT200UMT from Thorlabs, 200µm fiber core, NA 0.39) were implanted 0.3mm above the viral injections. For bilateral fiber implantations in DAT-Cre animals (both positive and WT) we observed an unusually high mortality of ∼30% during the first 24h after the surgery.

### Behavioral protocol

After a resting period of at least five days post-surgery, animals were habituated to handling and head restrain during a period of 3-4 days. Mice were head-restrained on a flat surface in a light-proof, sound-attenuated metal box. Their behavior was monitored using 2 cameras: a visible light camera placed on the side (to measure pupil dilation and facial movements; Mako-G040B, 30fps) and a far-infrared camera (FLIR A325sc, 50mm lens, 60 fps, 320x240 pixels, uncooled, microbolometer) placed below nostrils to non-invasively detect inhalations based on temperature changes, as described previously in (Mutlu et al., 2018).

#### a) Odor exposure

After the habituation period, all mice were exposed to odorants using a custom-made olfactometer (for detailed description, see Esquivelzeta Rabell et al., 2017) connected to a tube placed ∼2cm in front of animal’s nostrils. Each odor had a vapor concentration of 1Pa and was presented in a pseudo-random order for 2 seconds, with an inter-trial interval (ITI) drawn from an exponential distribution (mean = 20s, min = 2s, max = 30s). The mice were familiarized with a set of 4 or 6 odorants for 4 days (10 trials each, 1 session per day). Additionally, a ‘blank’ (mineral oil, which was used as medium for diluting odorants) was presented 10 times.

On the novelty day, all the familiar odorants were presented (8-10 trials each), accompanied by 6 novel odorants (8-10 trials each). The odors were presented in blocks ensuring that all of the odors would be presented at least once before moving to the next trial; the order of stimuli within a block was random. To gather more data (e.g., record electrophysiological activity from a different subset of Neuropixels channels), the novelty exposure was repeated 1-3 times, each day using a different set of 6 (previously not presented) odorants. For each animal, the odors used as novel and familiar were selected randomly from the following list of 24 chemicals: anisole, ethyl valerate, cinnamaldehyde, pentenoic acid, eugenol, limonene, trimethyl pyrazine, linalyl formate, geraniol, P.E.A, ethyl decanoate, nonadienal, benzaldehyde, heptanal, isoamyl acetate, octanoic acid, menthol, dimethylpyrazine, alfa methylstyrene, methyl anthranilate, alpha phellandrene, salicylaldehyde, thymol and geranyl acetate.

Because the optogenetic construct BIPOLES allowed to both activate or inhibit the same neurons (depending on light color), the same animals were used for both types of experiment. The inhibition was always performed on the 1^st^ novelty day: in addition to 6 novel and 6 familiar odors, 6 different novel odors were paired with blue light (‘novel+inhibition’ condition). During the next 3 consecutive days, all the odors not paired with light were presented 10 times. As a result, at this stage of experiment each animal was familiar with 12 different odorants. On the stimulation day (a.k.a. 2^nd^ novelty day) half of these odors were paired with red light (‘familiar+stim’ condition) and half not (‘familiar’ condition). Additionally, 6 truly novel stimuli were presented. Such design allowed us to test both effects of inhibition and stimulation with a limited number of odorants available. Odorants from all three categories (‘novel’, ‘familiar’ and ‘novel+inhibition’ or ‘familiar+stimulation’) were also presented the day after optogenetic manipulation, this time without any light, to test if long-term non-associative learning was affected (Suppl. Fig. 5).

#### b) Classical conditioning

The animals used for electrophysiological recordings from VTA/SNc, in addition to being exposed to odorants, underwent classical conditioning with rewards. Starting from the habituation period, their access to water was limited to 45 min/day. During the familiarization period, immediately after the odor exposure protocol was finished, a metal spout for water delivery was placed close to their mouth. Animals were exposed to 2 different cues, one followed by delivery of a big drop of water (∼4µL) and the other one by a small drop (∼1µL). The odorants used as cues were benzyl acetate and S+ carvone; their reward value was counterbalanced between the animals. During a training session, each odorant was presented 20 times for 2s; the reward was delivered with 100% probability 3.8s after the odor onset. The licking behavior was monitored using an optical distance sensor (Waveshare 9523), which was placed parrallel to the water spout and allowed to measure the distance between mouse tongue and the spout without creating electrical artifacts. The electrophysiological recordings from VTA/SNc were performed after either 4 or 8 days of the training (on 1^st^ or 2^nd^ novelty day, respectively). Because animals displayed clear anticipatory licking already after 4 days of training, data from both sessions is presented jointly (Fig. 1C).

### Data acquisition

#### a) Behavioral data

The experiment was controlled by a custom LabView software through an I/O board (PCIe-6351, National Instruments). Video recordings from both cameras were triggered 4s before valve opening and finished 8s after it; the ITI periods were not recorded. Videos from the face camera were saved as .tiff stacks using labcams software (https://github.com/jcouto/labcams). Videos from the thermal camera were saved as .seq files by ResearchIR Max software (v.4.40.9.30, FLIR). The National Instruments board collected all the digital (timing of odor delivery, timing of both cameras acquiring frames) and analog (readout from the optical distance sensor) signals with 1kHz sampling rate.

#### b) Electrophysiological data

The electrophysiological data was acquired continuously with a PXIe system (National Instruments), controlled by SpikeGLX software (https://github.com/billkarsh/SpikeGLX). The raw (broadband) data was collected with 30kHz sampling rate in a local reference configuration (reference from the joint shank tips).

#### c) Photometry data

Fiber photometry data was acquired using a commercial multi-fiber system (FP3002, Neurophotometrics), controlled by Bonsai software (https://bonsai-rx.org/). Briefly, the tissue was excited interchangeably with either 470nm or 415nm light at 80 fps rate (effectively, 40fps per channel). The emitted light was split into red and green channels with dichroic mirrors and collected by a CMOS camera. The light intensity at each region of interest (corresponding to individual fibers) was measured separately for the two colors and saved, while the raw images were discarded.

### Odor delay and synchronization

The olfactometer delivered odors with some delay, defined here as the period between valve opening and the volatile compound reaching the air in front of animal’s nose. Using photoionization detector (PID, 200B, Aurora Scientific), it was possible to determine this delay to be ∼600ms. In addition, the animals were able to perceive the odorant not immediately, but only when they performed the first inhalation after the 600ms delay period. To account for that, the neuronal responses to odorants were aligned to this time point (indicated as ‘Time from 1^st^ inhalation’ = 0) on the firing rate plots. The neuronal responses to rewards and optogenetic stimulation were aligned to the actual time of stimulus coming (irrespective of animal inhalations). For this reason, the raster plots demonstrating both cue and reward responses were created by concatenating two time windows separated by a varied delay, as illustrated by a dashed line on the time axis. To maintain consistency with the time of optogenetic stimulation, the behavioral responses were not re-aligned, and ‘Time from odor’ = 0 indicates the olfactometer valve opening.

### Optogenetics – stimulation protocol

For optogenetic stimulation, fiber-coupled LEDs (M455F3 or M625F2, Thorlabs) powered by an analogue driver (LEDD1B, Thorlabs) were used. The light was delivered to cannulas with a bifurcated (double-fiber) patch-cord (200µm core, 0.39NA, cat. # BFYL2LS01, Thorlabs). To avoid evoking artifacts in electrophysiological experiments, light was delivered as smooth sinusoidal patterns (see below) rather than binary pulses. In practice, an Arduino Uno Board was sending out pulse-width modulated (PWM) signals, smoothed by a simple resistor-capacitor circuit (R=1kΩ, C=10µF) before reaching the LED driver. For inhibition with blue light, the LED power was rapidly (∼30ms) increased to 100% at the beginning of odor delivery (i.e., valve opening), kept at maximum for 3 seconds, and then gradually ramped down to 0% over the course of 1 second. For stimulation with red light, a 0.5-second-long sinusoid (20Hz frequency) was delivered 100ms after the estimated time of odorant reaching the mouse (see section ‘Odor delay and synchronization’). The maximum estimated power (accounting for the light loss at the cannulas) was 5mW and 3.5mW for blue and red light, respectively. The same protocol was used for stimulating both VTA and brainstem.

### Histology

After behavioral experiments ended, animals were deeply anesthetized with pentobarbital (*Dolethal*), followed by transcranial perfusion with 0.1M phosphate-buffered saline (PBS) solution and 4% paraformaldehyde (PFA) solution. The brains were extracted and placed in PFA for 24 hours post fixation. After that, the brains were frozen and cut on cryostat in 50µm thick coronal slices. Free-floating sections were collected into 48 well plates filled with PBS. Sections from BIPOLES-injected animals were further processed for immunohistochemical staining according to the following protocol:

1. Wash slices in PBS (3 × 5 min).
2. Block for 1h in 10% NCS in PBST ([v/v], newborn calf serum [NCS] in 0.3% [v/v] Triton X-100 containing PBS) at room temperature.
3. Incubate for 40-48h at 4°C with primary antibody (anti-mCerulean polyclonal in goat, cat. #AB8208, Sicgen) at 1:250 in carrier solution (2% NCS in PBST)
4. Wash in PBS (3x 5-10 min).
5. Incubate for 3h at room temperature in carrier solution (same as above - 2% NCS in PBST) with secondary antibody at 1:1000 (Donkey anti-Goat, conjugated with Alexa Fluor 594, ThermoFisher/Invitrogen, cat. #A-11058).
6. Wash in PBS (3x 5-10 min).

All brain slices were mounted on glass microscopy slides with anti-fading medium (ProLong, cat. # P36934, Invitrogen). Afterward, all brain slices were imaged by either a confocal microscope (LSM 710 Zeiss) or slide scanner (SlideExpress 2, Nikon/Marzhauser).

### Atlas registration & exclusion criteria

The Neuropixels shanks and viral expression from the mdPons stimulation experiment were registered to the Common Coordinate Framework (CCF, Allen Brain Institute) using the HERBS software (Fuglstad et al., 2023). The positions of individual Neuropixels channels were adjusted to account for the CCF being stretched in DV direction (see https://community.brain-map.org/t/how-to-transform-ccf-x-y-z-coordinates-into-stereotactic-coordinates/1858). Finally, anatomical annotations for each voxel were obtained using the unified labeling framework (Chon et al., 2019) by matching the position in CCF to the nearest coronal slice in the unified labeling atlas (available in 100µm coronal resolution; https://kimlab.io/brain-map/atlas/).

Cells recorded outside of the target brain regions were excluded from further analysis. In the mdPons stimulation experiment, all the animals with viral expression in at least one hemisphere (n=7/8) were included. To create Fig. 4F, the data was combined by including from each animal the hemisphere with larger area labeled. In the DA inhibition/stimulation experiment, the animals were included only if viral expression and correct fiber placement could be verified for both hemispheres (n=8/14).

### Behavioral data analysis

To detect inhalation onsets, the videos from the nose camera were analyzed by a neural network (available upon request). The network was trained to reconstruct intranasal pressure signals measured invasively in a separate group of animals (Mutlu et al., 2018) based on sequences of thermal images. To plot sniffing rate after stimulus presentation, the obtained binary data (inhalation/no inhalation) was smoothed with a gaussian kernel (SD=0.15s, truncated at 4SDs), and in each trial subtracted from the breathing rate during baseline (-2 to 0s before olfactometer valve opening). All statistical comparisons were performed on unsmoothed sniffing rate calculated in a time bin (0.5 to 2.5s) and similarly subtracted from baseline (-2 to 0s).

For analyses relating neuronal activity to sniffing on a trial-by-trial basis, the sniffing response in each trial (Δ sniffing) was defined as the inhalation rate within 1 s after the first post-odor inhalation (i.e., with the same alignment as the neuronal responses), minus the breathing rate during the 1 s preceding valve opening in the same trial. Positive values therefore indicate sniffing above baseline. In the fiber photometry data, a 2-s response window and a 4-s baseline (the whole pre-odor period) were used, to account for the slower calcium dynamics.

To measure the amount of movement, face videos were analyzed using Facemap software (Syeda et al., 2024). The obtained signal reflected the total amount of changes in pixel values in the face region (after reducing the dimensionality of data using singular value decomposition, as described in (Stringer et al., 2019). Data was smoothed with a gaussian kernel (SD=0.15s, truncated at 4SDs), averaged in the same time bin as sniffing (0.5 to 2.5s) and normalized as % change relative to baseline. In the optogenetic stimulation experiments, the oscillating red light sometimes evoked artifacts in the calculated motion signal. For that reason, a time bin when light was no longer present (2 to 3s after valve opening) was used for statistical comparisons.

To measure pupil dilation, videos were converted from .tiff stacks to .avi and analyzed with a neural network (Mazziotti et al., 2021), using a Google Colab script (https://github.com/KacperKon/meye/blob/master/meye_colab.ipynb). Next, the data was post-processed: frames with high blink probability (>0.5) or pupil size differing extremely from the mean (+/-5 SD) were removed and interpolated based on neighboring values. Afterwards, the outliers were removed with a rolling median filter (size = 15 samples). The data was also smoothed with the same gaussian kernel as other behavioral signals and normalized as % change from baseline. Due to slower dynamics, time bins for pupil analysis were different than for sniffing: 1.5s to 3.5s (experiments with no red-light stimulation) and 2 to 3s (in experiments with possible red-light artifact).

For statistical comparisons, the mean value for each behavioral parameter was calculated using the first 4 presentations of all odorants from a given condition. In the combined inhibition-stimulation experiment, the mean response to novel and familiar odors was obtained for each mouse by calculating the average from 2 novelty days (as we have not observed systematic differences between them). For statistical testing, repeated-measures ANOVA was used, followed by post-hoc pairwise comparisons (paired t-tests with *p* values adjusted for multiple comparisons using the False Discovery Rate correction (Benjamini and Hochberg, 1995).

To determine the latency of the sniffing evoked by optogenetic stimulation, we used ZETA test implemented in Python (Montijn et al., 2021). Briefly, the method compared the cumulative number of inhalations after the stimulation onset to a control distribution, obtained by randomly shuffling the inhalations in time 50 times. The peak ZETA was defined as the moment when the observed values were maximally different from the control distribution (within the period 0-3 sec after the stimulation). As an indication of latency, the time needed to reach half of the peak was used. To ensure stable estimates, the results were obtained by repeating the whole procedure 100 times and averaging. Finally, the latencies were compared between the groups (VTA vs. mdPons stimulation) using t-test with Welch correction for unequal variances.

### Spike sorting and unit selection

Electrophysiological data were processed with an automated pipeline, available at https://github.com/KacperKon/SpikeSorting. First, the raw action-potential band was band-pass filtered (300– 10,000 Hz; 12th-order single-pass Butterworth filter, equivalent to a 3rd-order zero-phase filter), re-referenced to the global common average of all channels and corrected for transient artifacts using CatGT (https://billkarsh.github.io/SpikeGLX). Spikes were then detected and sorted with Kilosort 4 (v4.1.7; Pachitariu et al., 2024) using default parameters (universal and learned template thresholds of 9 and 8, rigid drift correction). Spike times were aligned to the clock of the behavioral acquisition system with TPrime, using a 1-Hz synchronization pulse recorded by both systems. For each sorted unit, waveforms (500 randomly selected spikes), templates and standard quality and waveform metrics were computed with SpikeInterface (v0.103.2; Buccino et al., 2020). Units were then classified with UnitRefine (Jain et al., 2025), a machine-learning classifier pretrained on manually curated mouse Neuropixels recordings (model “UnitRefine-mice-sua-classifier”), and only units classified as single units were included; the classifier output was inspected visually but not modified. Finally, units that did not fire a single spike in more than half of the trials (presence ratio < 0.5) were excluded. AON data were taken from a previously published dataset (de Plus et al., 2025). Preprocessing, spike sorting and unit selection were performed in the original study, and the resulting single units were used here without re-sorting.

### Firing rates

If not stated otherwise, firing rates were calculated in 100-ms bins. Odor responses were aligned to the first inhalation after odor arrival (see “Odor delay and synchronization”), and the neuronal response in each trial was defined as the mean firing rate 0–500 ms after this inhalation. Where z-scored responses are reported, the firing rate in each trial was z-scored relative to the baseline of the same trial (2 s before the first inhalation; bins with a baseline SD below 0.1 spikes/s were scaled by 1 instead). The grand average response to novel and familiar odors (not split by active vs. passive condition) was calculated from the first 4 presentations of each odorant. To determine whether a cell population responded more strongly to novel than to familiar stimuli, mean firing rates (0–500 ms) were compared with a Wilcoxon signed-rank test (n = number of cells). To test whether individual cells responded to novel stimuli, responses (0–500 ms after the first inhalation) were compared with the baseline (2 s before) using a Wilcoxon signed-rank test (n = number of trials); p values were FDR-corrected separately for each cell population, and cells were classified as responsive if q (adjusted p) < 0.05.

### Cell type classification

To cluster cells according to their putative type, we used the response properties described in the optotagging study by (Cohen et al., 2012). First, firing rates from big reward trials were normalized by calculating auROC relative to baseline period (for details, see (Cohen et al., 2012). This procedure allows to quantify how reliably each time bin can be discriminated from baseline, based on the distribution of firing rates across trials. The auROC score of 1 indicates that in all trials firing rate at given moment was higher than all values in the baseline distribution; score of 0 that in all trials they were lower; score of 0.5 indicates no difference from baseline.

Next, cells were classified into four categories based on their auROC values (100-ms bins). For each cell, the auROC value deviating most from 0.5 was determined within the cue window (0–0.6 s after the first inhalation) and the reward window (0.1 s before to 0.5 s after reward delivery). Cells for which this value remained between 0.4 and 0.6 were categorized as non-responsive (Non-resp), cells with values below 0.4 as inhibited (Inh.), and cells with values above 0.6 as responsive. Responsive cells were further categorized based on their activity during the delay period (-1.1 to -0.1s) before reward delivery: cells with a mean auROC above 0.56 were classified as pGABA, and the remaining cells as pDA. Only cells located in the VTA, SNc or RRF were classified. The classification was further verified by plotting the distribution of baseline firing rates and spike widths for each cell type (Suppl. Fig. 1).

### Active vs. passive trials and spontaneous exploration

All novel and familiar odor presentations (excluding blank, reward and optogenetic stimulation trials) were classified as “active” (Δ sniffing > 0) or “passive” (Δ sniffing ≤ 0). Because novel odors evoked more sniffing than familiar odors, trial sets were then matched within each session, so that novel and familiar trials had the same mean Δ sniffing within each sniffing level (Suppl. Fig. 2). Among active trials, trials were removed from the condition with the higher mean, starting from the highest Δ sniffing, until its mean reached that of the other condition. Among passive trials, trials were removed from the condition with the lower mean, starting from the lowest Δ sniffing, until its mean reached that of the other condition. Removal stopped before the mean crossed the target value; a match was accepted if the remaining difference between the condition means was ≤ 0.25 inhalations/s, and otherwise the corresponding sniffing level was not analyzed for that session.

Spontaneous episodes of active exploration (sniffing bouts) were detected in the pre-odor period of each trial (excluding reward trials). A bout was defined as a sequence of at least 6 consecutive inhalations with all inter-inhalation intervals shorter than 1/3.5 s (instantaneous rate > 3.5 Hz), with its onset at the first inhalation. Only bouts starting between 2 s before and the moment of the olfactometer valve opening were included, so that a baseline period preceded each bout. The Δ sniffing of a bout was calculated as the inhalation rate during its first 0.5 s minus the baseline breathing rate of the same trial (1 s before valve opening, as for odor trials). Bouts with Δ sniffing ≤ 0 were discarded, and the remaining bouts were matched to the mean Δ sniffing of the active odor trials of the same session, using the same procedure as above (removing bouts with the highest Δ sniffing).

Firing rates around bout onsets were calculated in 100-ms bins (from 2 s before to 3 s after onset) and z-scored relative to the baseline from 2 s to 0.5 s before bout onset; the last 0.5 s before onset was excluded from the baseline because neuronal activity often started to increase before the first inhalation of a bout. For each neuron, z-scored responses were averaged across the matched trials of five conditions: novel-active, familiar-active, novel-passive, familiar-passive and spontaneous bouts. Fiber photometry data (single-fiber examples, Fig. 3F) were analyzed in the same way, with two differences. First, the z-scored ΔF/F signal (see “Photometry data analysis”) was only baseline-subtracted, using the mean from 1.5 to 0.5 s before bout onset. Second, because photometry was recorded only during trials, bouts were detected from trial onset up to valve opening, and bouts whose baseline did not lie within the recorded trial were excluded.

Statistics. For each population, the mean z-scored responses of each neuron (0–500 ms after the first inhalation) in the four odor conditions were analyzed with linear mixed models (statsmodels), with novelty, sniffing level (active vs. passive) and their interaction as fixed effects, and random intercepts for session and for neuron nested within session. Each effect was tested with a likelihood-ratio test comparing maximum-likelihood fits of the models with and without the term (main effects against the main-effects model, the interaction against the full model). Novel and familiar responses were compared post hoc within active and within passive trials (separate mixed models, Wald tests, FDR-corrected). Differences between populations (mdPons vs. SC) were tested with a combined model additionally including population and its interactions. Responses to spontaneous bouts are shown for comparison but were not included in the models.

### Population trajectories

For each population, neurons with data in all five conditions were included. The condition-averaged z-scored responses from 0.5 s before to 2 s after the first inhalation (or bout onset; 100-ms bins) were arranged into a matrix of time points × neurons, the five conditions were concatenated along the time axis, and principal component analysis was fitted to this matrix (10 components). Population trajectories (Fig. 2D) show the projections of each condition onto the first two components; for display only, trajectories were smoothed with a causal half-Gaussian kernel (SD = 200 ms).

Distances between conditions were computed in the full neuronal space (without dimensionality reduction): for each time bin from 0 to 2 s, the Euclidean distance between the population vectors of two conditions was calculated, averaged across time bins, and normalized by the square root of the total variance of the population activity (summed across neurons, over all time points and conditions). Three contrasts were computed: novel vs. familiar (average of the active and passive pairs), active vs. passive (average of the novel and familiar pairs) and spontaneous vs. odor-evoked exploration (average of spontaneous vs. novel-active and spontaneous vs. familiar-active). To estimate the distance expected by chance, the trial labels of the two conditions in each pair were shuffled within each session (one permutation per session, shared by all its neurons) and the condition means and distances were recomputed (200 permutations). Distances in Fig. 2E are shown relative to this null distribution (observed distance minus the mean of the null distribution). Statistical significance was assessed with a permutation test, p = (1 + number of permutations with a distance ≥ the observed one) / (number of permutations + 1), FDR-corrected across all populations and contrasts. Standard errors were estimated as the standard deviation of the distances recomputed on neurons from random subsets of 50% of the mice (500 subsamples, drawn without replacement).

### Single cell response properties

To quantify cross-correlations between neuronal and behavioral activity, instantaneous firing rate was calculated for each cell using the same gaussian kernel (SD=0.15s, truncated at 4SDs) as the one used for smoothing behavioral signals (see above). The data from all novel and familiar trials (without optogenetic manipulation) was concatenated and the cross-correlations were computed using the ‘scipy.signal’ package with 30 samples/second resolution. Face camera data from some sessions was lost due to experimenter’s error (3/26 VTA/SNc recordings) and could not be included in the calculations.

To verify if the correlation with sniffing was present in all conditions, a Pearson correlation (lag=0) was calculated separately for baseline periods (-3.5s to 0s before valve opening) and odor periods (novel or familiar, first 4 presentations, 0 to 5.5s).

To measure how strongly each cell was modulated by each behavioral variable (Fig. 3B), cross-correlations between firing rate and behavior were computed using baseline (pre-odor) periods only (3.5 s before valve opening, all novel and familiar trials). Because baseline segments were short, trials were not concatenated; instead, for each lag the correlation was computed from pairs of samples belonging to the same trial (with both signals z-scored across all baseline samples). The modulation was defined as the cross-correlation value at the lag with the largest absolute value within ±1 s. The odor response of each neuron was its mean z-scored response 0–500 ms after the first inhalation, averaged across novel and familiar trials. Novelty selectivity was the difference between novel and familiar responses in the same window, computed on the sniffing-matched trial sets (see above) separately for active and passive trials and then averaged. The reward response was the mean z-scored response from 0.1 s before to 0.5 s after big-reward delivery, and the response to optogenetic stimulation was calculated in the first 100 ms after laser onset (no-odor stimulation trials). Spearman correlations were computed between all these metrics (together with the mean baseline firing rate), p values were FDR-corrected (Benjamini and Hochberg, 1995), and only correlations with q < 0.05 are shown.

### Regression

Ordinary least-squares linear regression was used to quantify how strongly the responses of each neuron were influenced by sniffing and stimulus novelty. The dependent variable was the response in each trial (mean firing rate 0–500 ms after the first inhalation, minus the mean firing rate during the 2 s before it). All novel and familiar odor presentations were included (excluding blank, reward and optogenetic trials). The model included an intercept and the following predictors: Δ sniffing (see “Single-trial sniffing response”), odor novelty (novel = 1, familiar = 0) and the novelty × Δ sniffing interaction.

Because the predictors were correlated (sniffing was higher for novel than for familiar odors), beta values could not be used to estimate the contributions of individual variables. Instead, the unique contribution of each variable was estimated with a permutation approach: the predictor(s) of interest were randomly shuffled across trials (100 permutations), the full model was refitted, and the unique contribution was defined as the mean decrease in R² relative to the unshuffled model. Three predictor groups were tested: sniffing (Δ sniffing and the interaction term), novelty (novelty and the interaction term) and the interaction term alone. Overall model significance was tested in the same way by shuffling all predictors jointly. For each neuron, the p value was the proportion of permutations in which R² exceeded that of the unshuffled model, and p values were FDR-corrected across neurons, separately for each predictor group. Neurons were classified as regulated by sniffing, by novelty or by both based on the significance of the respective unique contributions (Fig. 3E).

At the population level, unique contributions were compared with a mixed-design ANOVA (between-subject factor: cell type or recorded region; within-subject factor: predictor group, using Python package pingouin), with Greenhouse–Geisser correction where sphericity was violated, followed by pairwise t-tests (FDR-corrected). The total variance explained by the full model was compared between populations with a Mann– Whitney U test. The same analysis was repeated in a short window just after odor delivery (0–100 ms; Suppl. Fig. 2). For fiber photometry, the dependent variable was the mean z-scored ΔF/F 0–2 s after the first inhalation (minus the 1 s before it), 1,000 permutations were used, and predictor groups were compared with a repeated-measures ANOVA.

### SVM

A linear support vector machine (SVM; scikit-learn LinearSVC) was trained to discriminate between two trial categories based on the responses of all recorded cells of a given type, pooled across sessions into a pseudo-population. The response of each cell in each trial was its mean firing rate 0–500 ms after the first inhalation minus the mean firing rate during the 2 s before. In each iteration, 36 trials (18 per category) were drawn from each session and combined across sessions into pseudo-trials; 26 trials (13 per category) were used for training and 10 for testing. Each cell’s responses were z-scored using the mean and SD of the training set only. The analysis was run in three conditions. In the first, novel and familiar trials were selected randomly and the SVM decoded novelty. In the second, novel and familiar trials were pair-matched for identical Δ sniffing, and the SVM again decoded novelty. In the third, trials were split into high (Δ sniffing above the session mean) and low sniffing (at or below the mean), with equal numbers of novel and familiar trials in each group, and the SVM decoded high vs. low sniffing. The procedure was repeated 300 times per condition. Decoding accuracies were compared with a mixed-design ANOVA (between-subject factor: population; within-subject factor: decoding condition), followed by pairwise t-tests (FDR-corrected), with resampling iterations as observations.

### Photometry data analysis

The multi-fiber photometry data (n=9 mice, 12 channels each) was preprocessed using the Python pipeline provided by (Martianova et al., 2019). The preprocessing included: 1) smoothing data with a flat window (length = 20 samples), 2) standardizing it as zdF/F and 3) de-trending and correcting for calcium-independent fluorescence changes by regressing out the contribution of red (isosbestic) from green (calcium-related) channel. The ‘bad’ channels (in which fluorescence reflected noise rather than neuronal activity) were identified manually and removed from the analysis. Due to the size of array being much bigger than the size of dopaminergic nuclei, most channels were excluded, and the final dataset consisted of 35 channels from n=8 mice (1/9 excluded due to no detectable signals).

## Acknowledgements

This work was supported by Interne Fondsen KU Leuven C14/21/111 (SH), Fonds Wetenschappelijk Onderzoek-Vlaanderen (FWO-Flanders) fellowship 1276122N (KK) and grant G097022N (SH).

## Author Contributions

KK and SH designed experiments and wrote the manuscript; KK, XT, SL and LDP performed the experiments; KK analyzed the data.

