## Supplementary Materials for "Active sensory exploration is a major driver of dopamine neuron activity"

| Ref. | Region | Response to sensory stimuli | Rapid habituation | Modality | Species | Reward present | Latency | Observed behaviors |
|---|---|---|---|---|---|---|---|---|
| Lak et al., 2016 | not reported | **yes** | **yes** | visual | macaque | yes | <200ms | not reported |
| Solie et al., 2022 | VTA | **yes** | **yes** | multi-modal | mouse | no | not reported | social inter., sniffing |
| Steinfels et al., 1983 | SN | **yes** | **yes** **no** | visual & auditory | cat | no | <70ms | orienting not reported |
| Ljungberg et al., 1992 | VTA / SN | **yes** | **no** | multi-modal | macaque | yes | <200ms | saccadic eye movements |
| Horvitz et al., 1997 | VTA | **yes** | **no** | visual auditory | cat | no | <70ms <50ms | not reported |
| Takeuchi et al., 2016 | VTA | **yes** | **no** | multi-modal | mouse | no | not reported | not reported |
| Kamiński et al., 2018 | SN | **yes** | **no** | visual | human | no | <500ms | button press |
| Ogasawara et al., 2022 | SN | **no** | **no** | visual | macaque | no | no responses | saccadic eye movements |
| Freeman et al., 1986 | VTA | **yes** | not reported | auditory & tactile | rat | no | not reported | orienting, sniffing |
| Mikell et al., 2014 | SN | **yes** | not reported | auditory | human | no | <300ms | not reported |

**Supplementary Table 1.** Summary existing of findings on midbrain dopamine neurons responses to novel stimuli. Only studies that reported activity of individual cells were included.

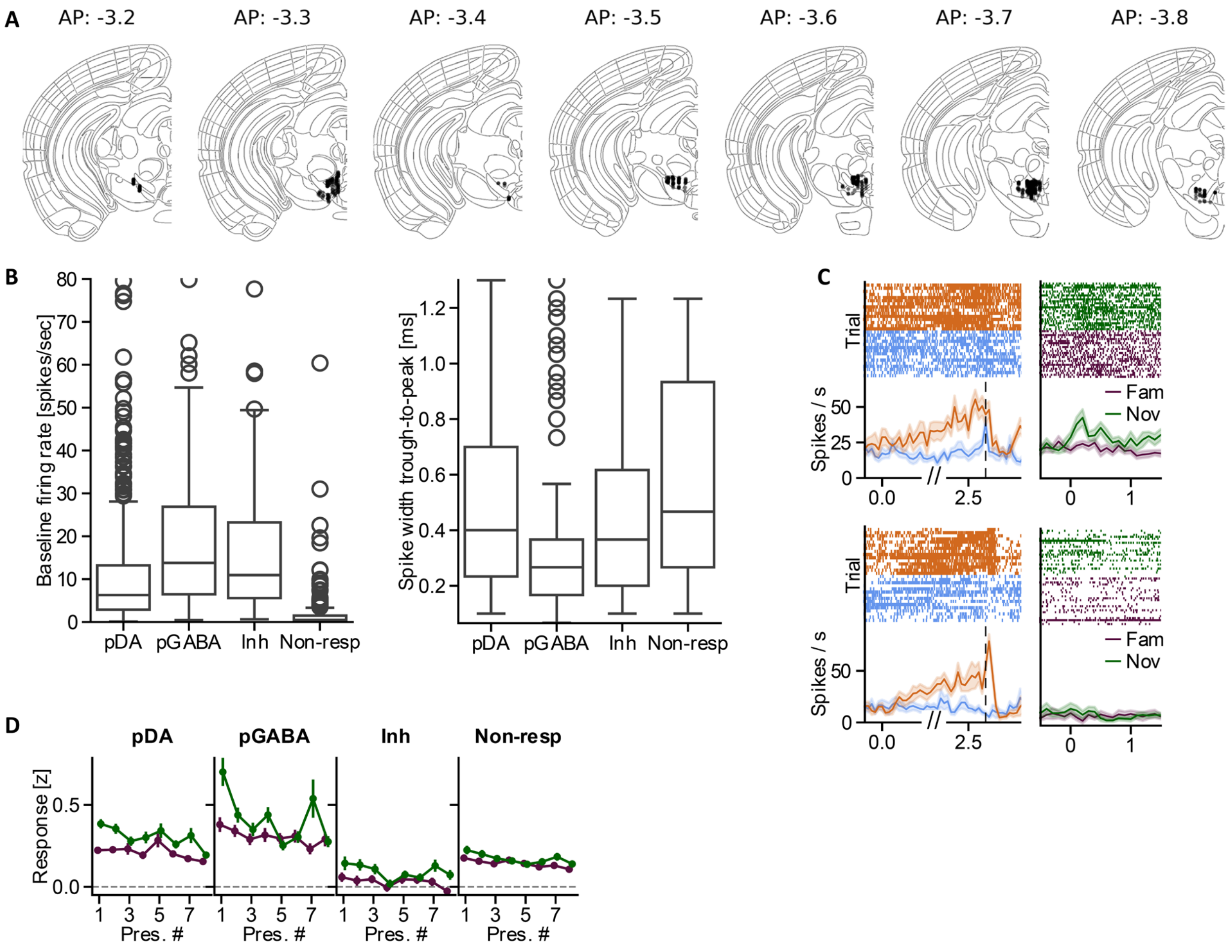


**Supplementary Figure 1.** A) Location of all the cells recorded from VTA, SNc or RRF. The numbers above each atlas plate correspond to anterior-posterior coordinates (in mm relative to bregma). B) Baseline firing rate (left) and spike width (right) of all cells recorded from VTA/SNc/RRF during the reward session, split into putative type. C) Example responses of two pGABA neurons, one with strong responses to novel stimuli (top) and one with weak ones (bottom). Orange: big reward, blue: small reward, green: novel, purple: familiar. Vertical line: presentation of reward. D) Habituation of neuronal responses across repeated stimulus presentations, showed separately for novel (green) and familiar (purple) odors.

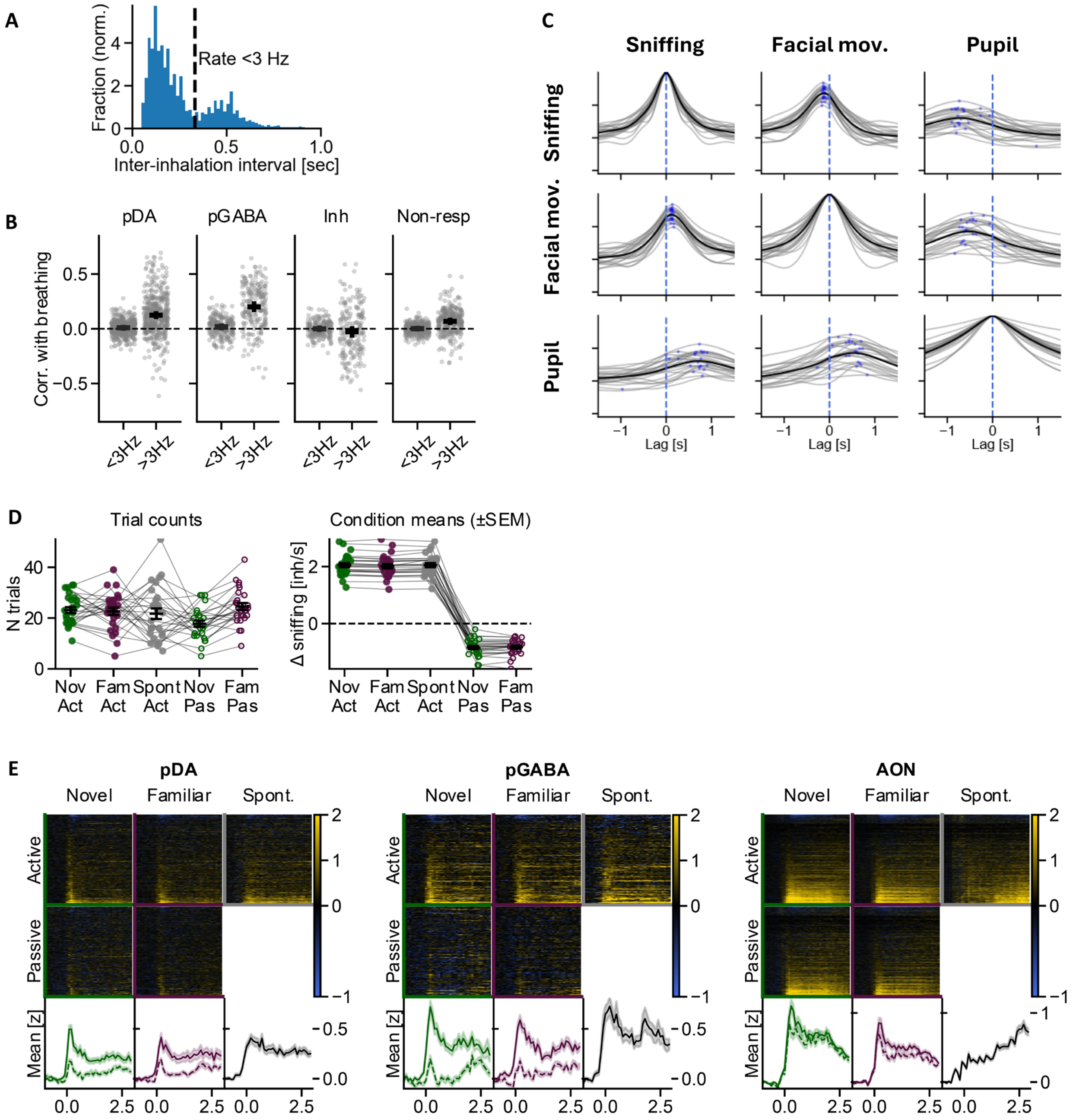


**Supplementary Figure 2.** A) Example distribution of inter-inhalations intervals from a representative mouse. Vertical line: interval corresponding to breathing at exactly 3Hz rate. B) Correlation between spiking and instantaneous breathing rate, calculated separately for slow breathing (<3Hz) vs. sniffing (>3Hz). C) Cross-correlations between the recorded behavioral signals. Thin gray lines: individual experimental sessions (n=23 from 13 mice); black lines: average; blue dots: peak cross-correlation in each session. D) Diagnostics of the trial-matching procedure, used to split trials into Active vs. Passive categories (see Methods & Fig. 2). Left – number of trials used per experimental session, right – the resulting mean sniffing response across conditions. Individual dots represent individual behacioral sessions. Nov – novel, Fam – familiar, Act – active, Pas – passive, Spont – spontaneous. E) The data from figure 2C, plotted together with the heatmaps illustrating responses of the single units (top panels).

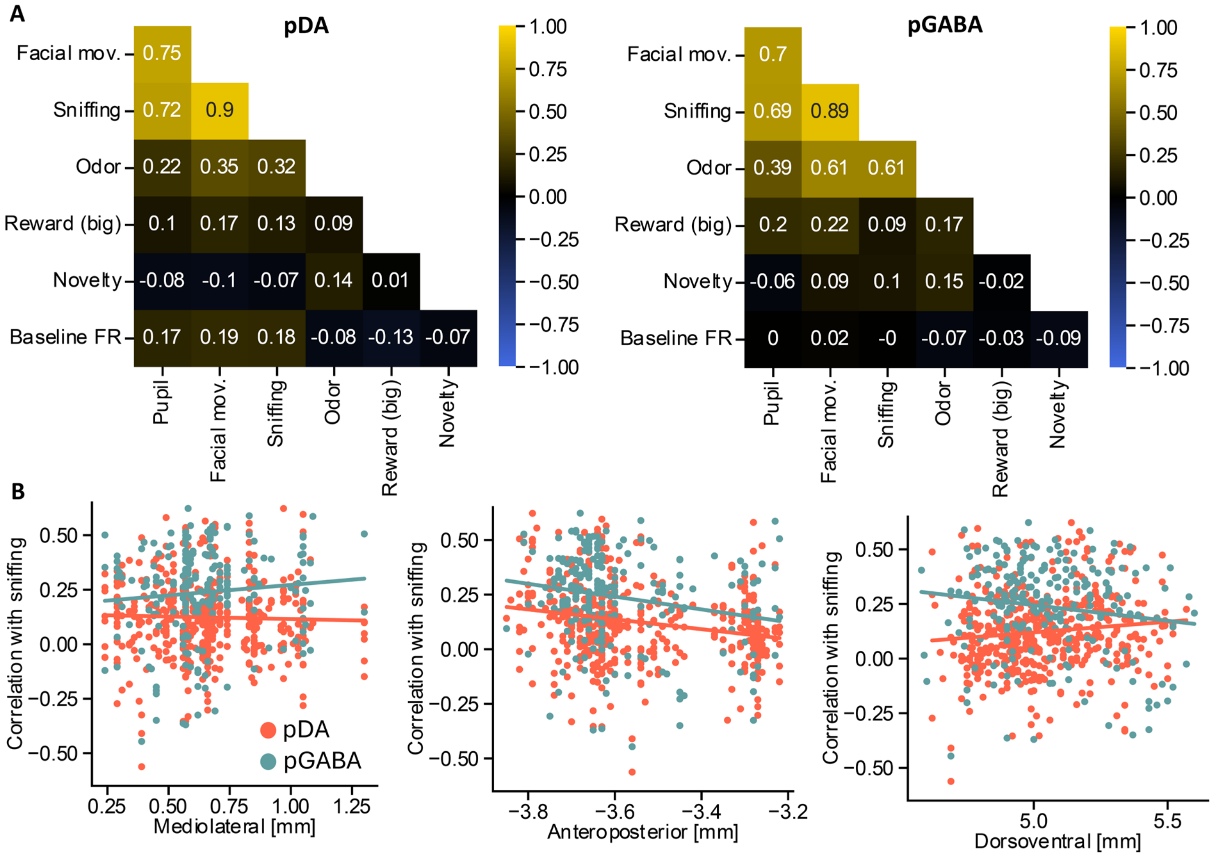


**Supplementary Figure 3.** A) Correlation matrix between different modulators of neuronal activity, plotted separately for pDA (left) and pGABA cells (right). B) Spearman's correlation between the degree of modulation by sniffing and anatomical location of pDA (red) and pGABA (green) cells. Left: mediolateral; pDA: r=−0.05, p>0.05; pGABA: r=0.09, p>0.05. Middle: anteroposterior; pDA: r=−0.26, p<0.001; pGABA: r=−0.19, p<0.01. Right: dorsoventral; pDA: r=0.12, p<0.01; pGABA: r=−0.10, p>0.05. The p values were not adjusted for multiple comparisons.

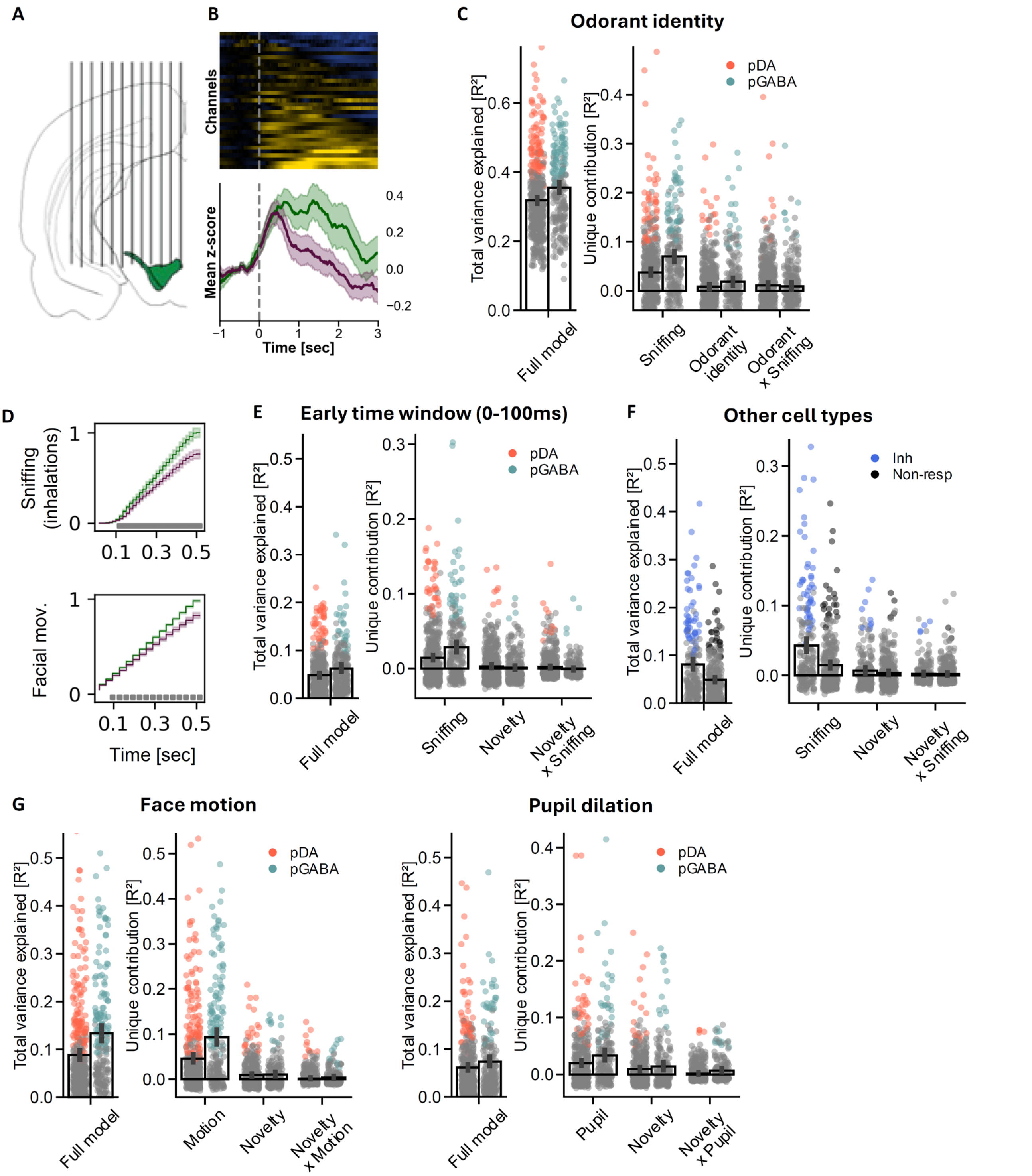


**Supplementary Figure 4.** A) Fiber placement in the photometry experiment. A linear array of 12 thin (100µm) optic fibers was used. B) DA responses to odors recorded with fiber photometry. Top: average responses from all the fibers to novel stimuli (n=35 channels with calcium signal from n=8 mice). Bottom: average responses to novel (green) vs. familiar (purple) stimuli. C) Regression analysis from Fig. 3D, repeated with chemical identity included as a predictor. D) Cumulative changes in behavior during the first 0.5 seconds after the presentation of novel (green) vs. familiar (purple) stimuli, aligned to the first inhalation. Top: sniffing, bottom: facial movements. Gray rectangles on the bottom indicate time points in which behavior was significantly different between novel vs. familiar condition ($p<0.05$, FDR adj.). E) Regression analysis from Fig. 3D, repeated on a smaller time window (0-100ms after the 1$^{st}$ inhalation). F) Regression analysis from Fig. 3D, repeated on the remaining cell types (inhibited and non-responsive to rewards). G) Regression analysis from Fig. 3D, repeated using facial motion (left) and pupil dilation (right) as indicators of active exploration.

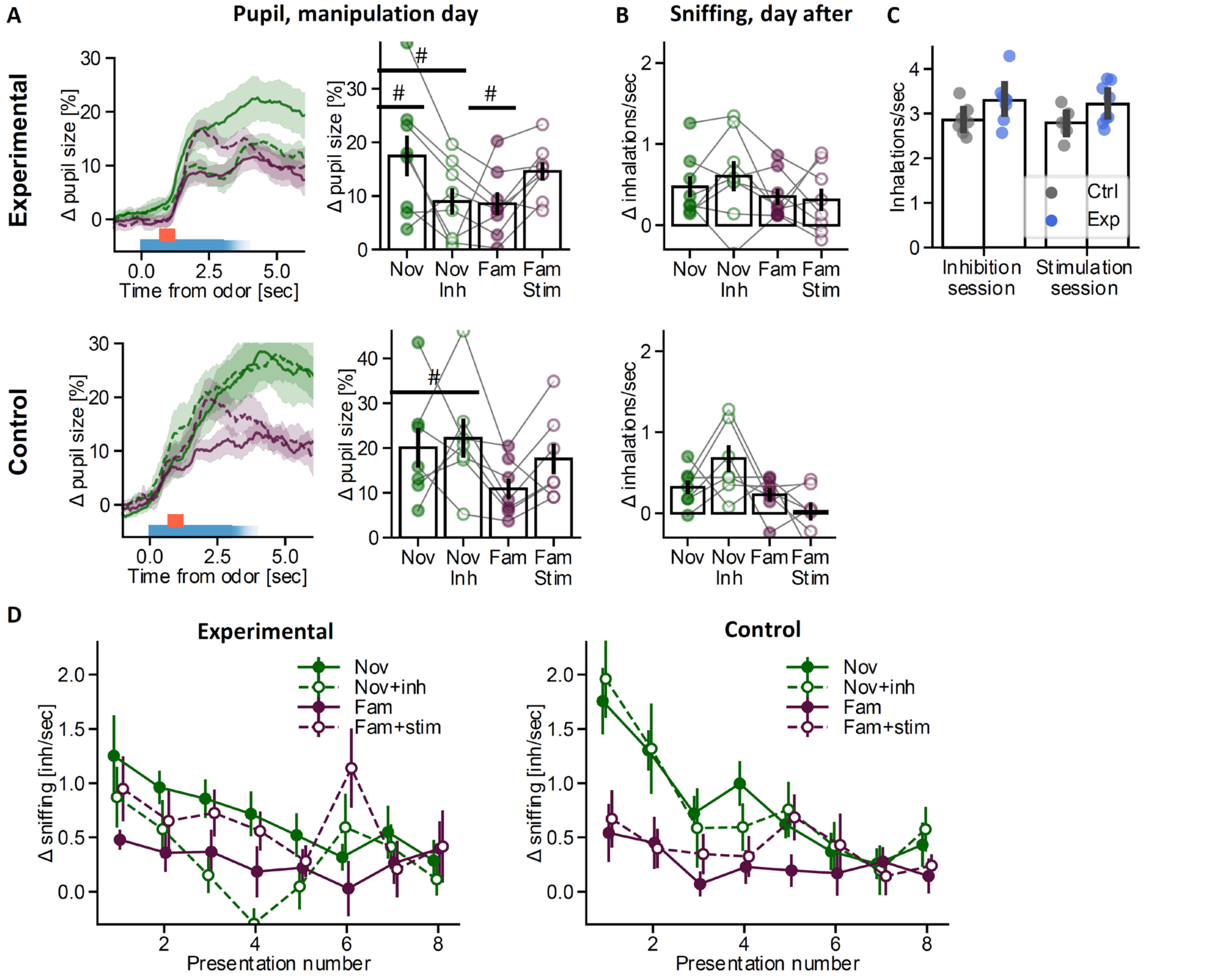


**Supplementary Figure 5.** A) The effects of DA manipulation (see Fig. 4) on pupil size. There was a trend for decreasing pupil dilation after DA inhibition in the experimental group (t(7)=-2.69, p=0.064), but not in the control group (t(6)=0.36, p=0.73). Similarly, stimulating DA neurons was associated with a trend to increase pupil size in the experimental group (t(7)=3.57, p=0.055), but not in the control group (t(6)=2.33, p=0.12). All p values were adjusted for multiple comparisons with FDR correction (see Methods). B) The effects of the optogenetic manipulation on long-term habituation. When tested next day after the manipulation, there were no differences in how animals responded with sniffing to odors paired vs. not paired with light. The results indicate that inhibiting DA activity does not prevent long-term habituation to novel odors. C) The comparison of baseline breathing rate between the groups. The values were obtained by averaging the breathing rate during baseline (4 sec before odor presentation) across all the trials used for the statistical analysis (first 4 presentations of all odors). The baseline breathing rate did not differ significantly between the groups neither in the inhibition (t(13)=1.98, p=0.07) nor the stimulation session (t(13)=1.91, p=0.08). The p values were not adjusted for multiple comparisons. D) Habituation curves: average sniffing in the DA manipulation experiment, plotted across odor presentations. Left: experimental group, right: control group.

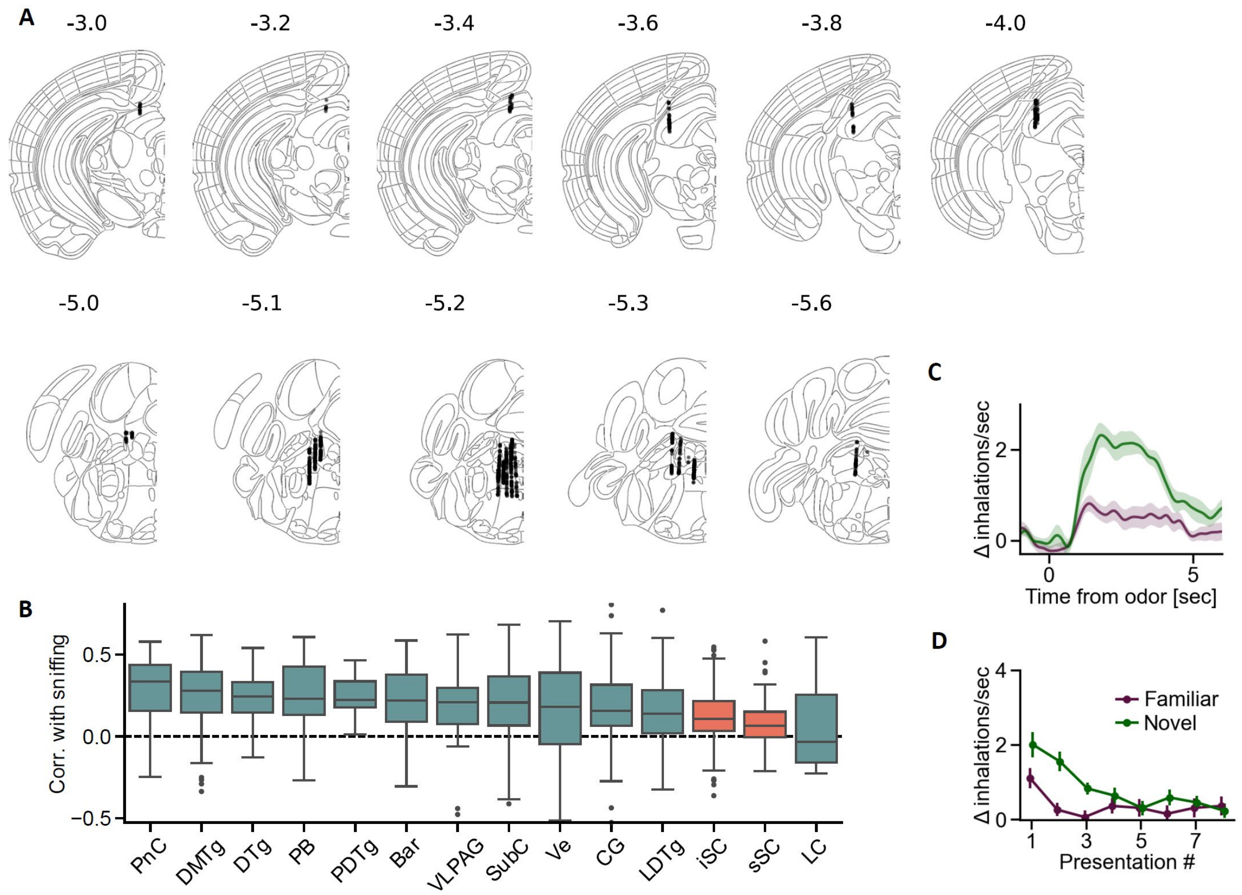


**Supplementary Figure 6.** A) Location of all the cells recorded from SC (top) or mdPons (bottom). The numbers above each atlas plate correspond to anterior-posterior coordinates (in mm relative to bregma). B) Distribution of correlations between spiking rate and sniffing rate (during pre-odor periods), plotted individually for all individual brain structures (mdPons, blue; SC, red). The structures are ordered by median correlation, from highest to lowest. PnC – pontine reticular nucleus, caudal part (n = 31); DMTg – dorsomedial tegmental area (n = 102); DTg – dorsal tegmental nucleus (n = 35); PB – parabrachial complex (n = 38); PDTg – posterodorsal tegmental nucleus (n = 8); Bar – Barrington's nucleus (n = 72); VLPAG – ventrolateral periaqueductal gray (n = 30); SubC – subcoeruleus nucleus (n = 130); Ve – medial vestibular nucleus (n = 39); CG – central gray (n = 157); LDTg – laterodorsal tegmental nucleus (n = 162); iSC – intermediate layers of SC (n = 117); sSC – superficial layers of SC (n = 55); LC – locus coeruleus (n = 6). C) Average sniffing responses in sessions in which SC or mdPons neurons were recorded (n=12 sessions from 6 mice). D) Average habituation curve from the same sessions.

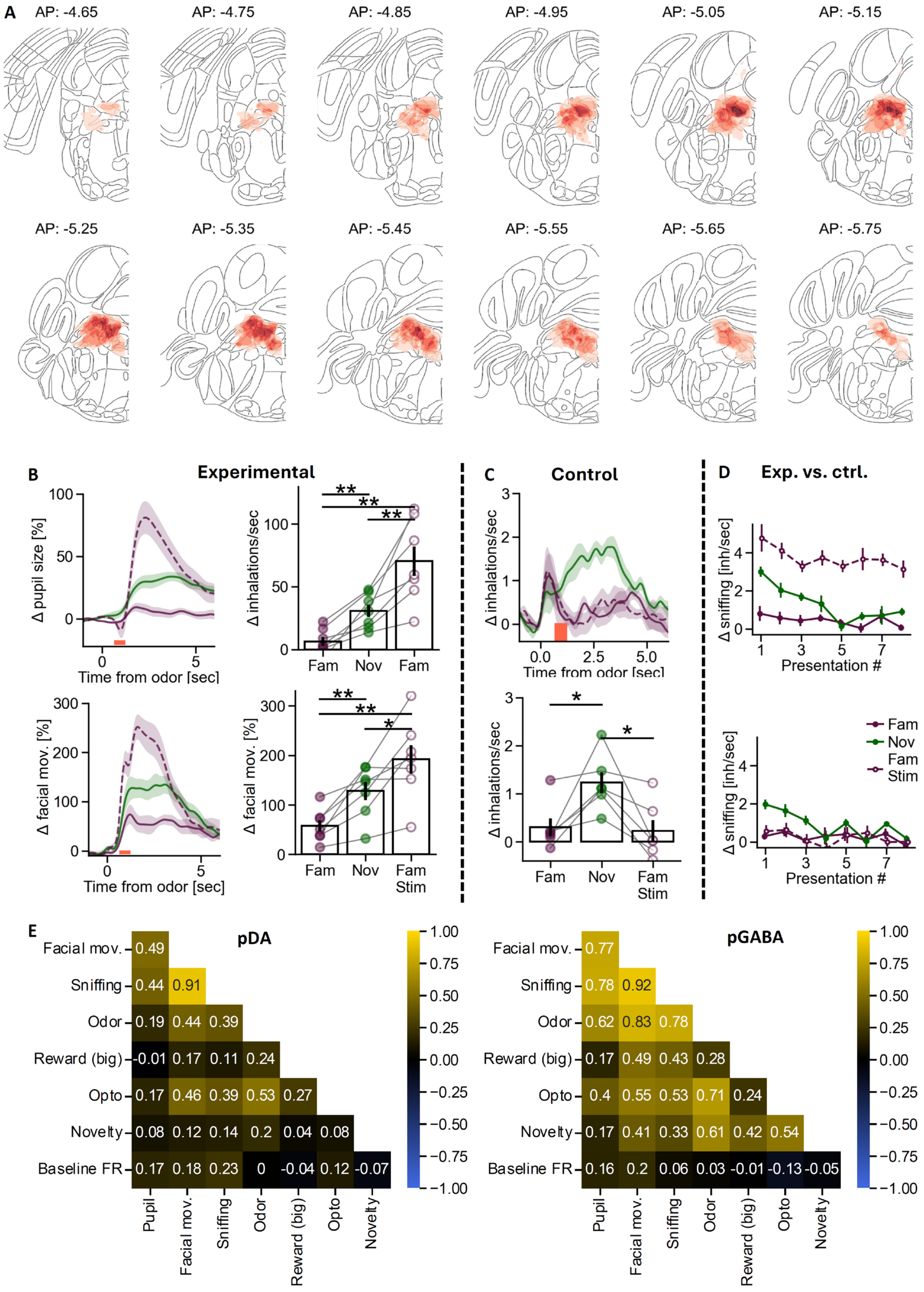


**Supplementary Figure 7.** A) Expression of the opsin (som-BIPOLES) in VGlut2 cells of mdPons across all animals (n=7). The color scale on the right indicates in how many animals (from 0 to 7) the virus was expressed in each location. The list of structures labelled in all animals included (ordered by % coverage): lateral parabrachial nucleus, laterodorsal tegmental nucleus, mesencephalic trigeminal tract, barrington nucleus, locus coeruleus, superior medullary velum. B) The effects of VGlut2 mdPons stimulation (see Fig. 5F & 5J) on pupil dilation (top) and facial movements (bottom). C) The effects of VGlut2 mdPons stimulation in the control group. D) Habituation curves: sniffing across odors presentations in the experimental group (top) vs. control group (bottom). E) Correlation matrix between different modulators of VTA/SNc neuronal activity in the VGlut2 mdPons stimulation experiment (see Fig. 5F & 5J). Left: pDA cells (n=78), right: pGABA cells (n=20).

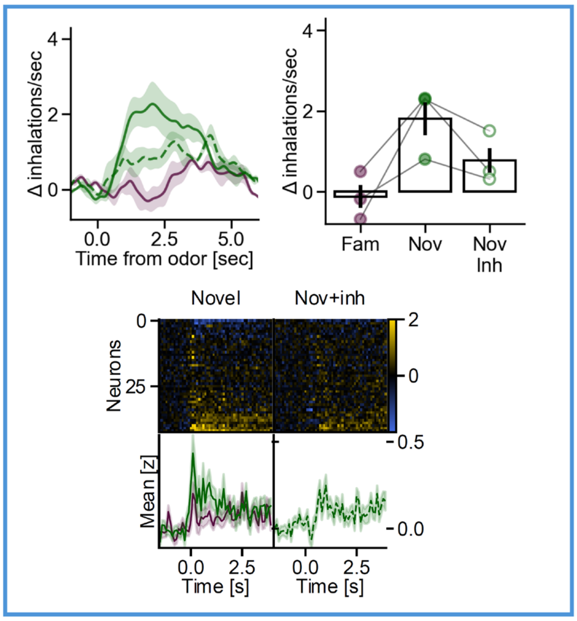
Δ inhalations/sec
Time from odor [sec]
Fam
Nov
Nov Inh
Novel
Nov+inh
Neurons
Mean [z]
Time [s]

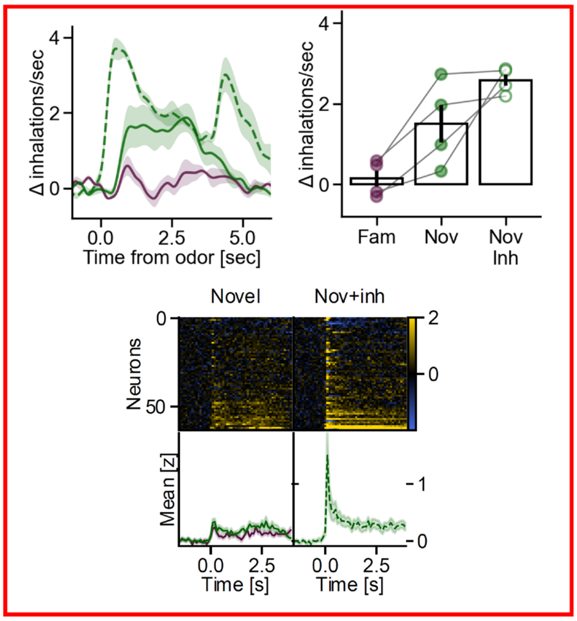
Δ inhalations/sec
Time from odor [sec]
Fam
Nov
Nov Inh
Novel
Nov+inh
Neurons
Mean [z]
Time [s]

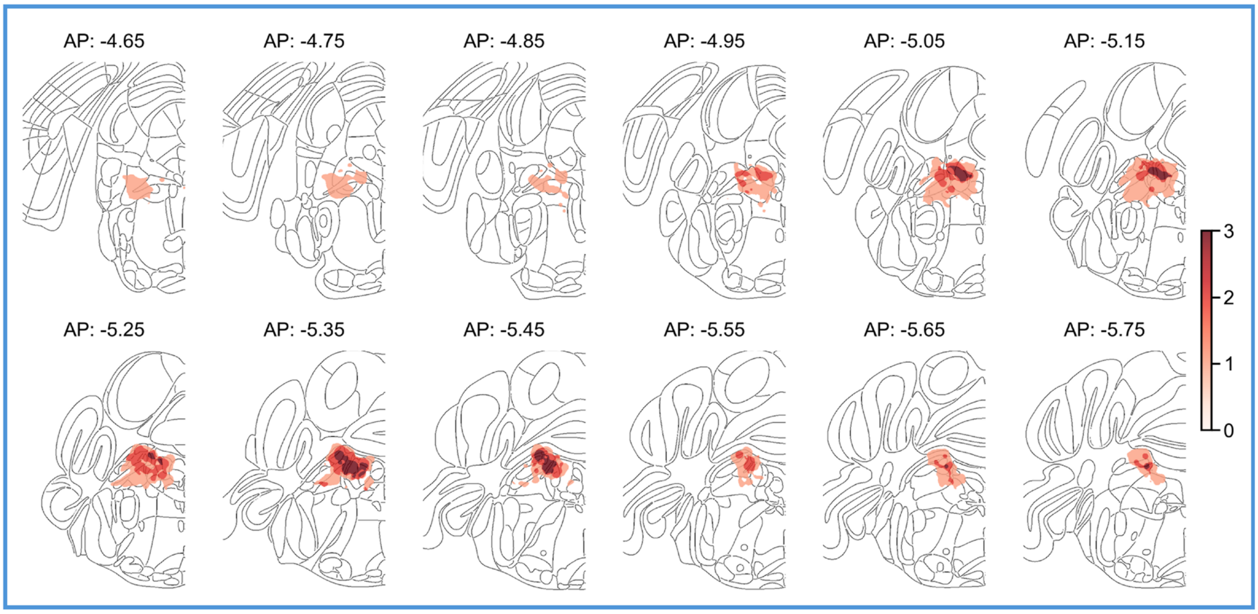
AP: -4.65
AP: -4.75
AP: -4.85
AP: -4.95
AP: -5.05
AP: -5.15
AP: -5.25
AP: -5.35
AP: -5.45
AP: -5.55
AP: -5.65
AP: -5.75

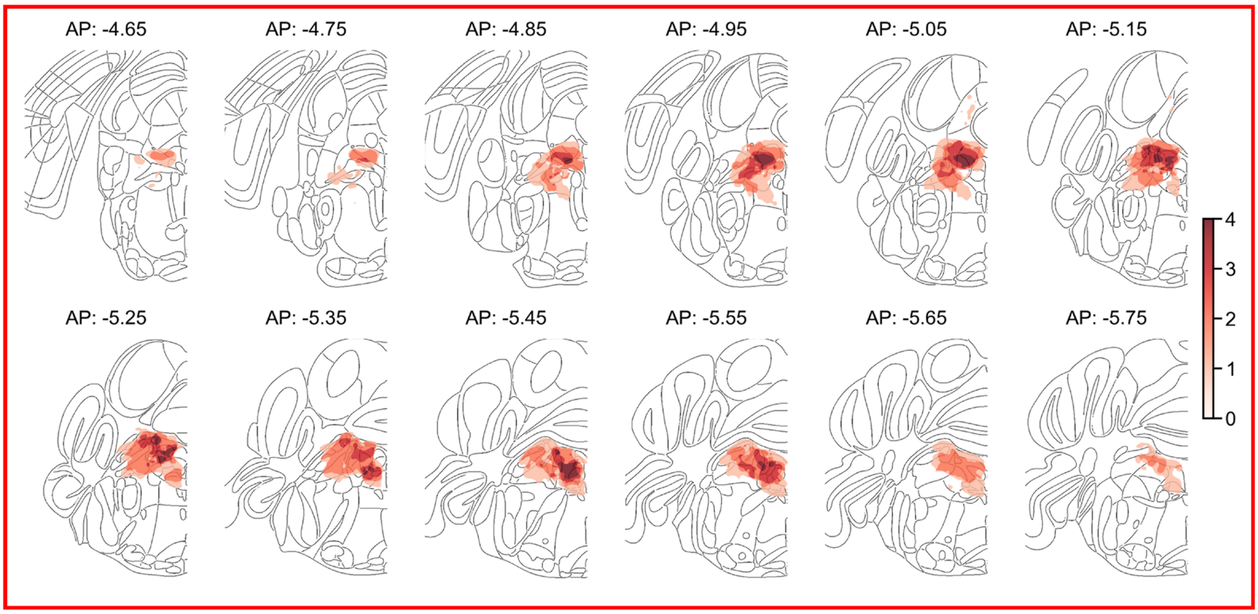
AP: -4.65
AP: -4.75
AP: -4.85
AP: -4.95
AP: -5.05
AP: -5.15
AP: -5.25
AP: -5.35
AP: -5.45
AP: -5.55
AP: -5.65
AP: -5.75

**Supplementary Figure 8.** The effects of inhibiting VGlut2 mdPons neurons with blue light. A) Blue frame: in 3 animals, inhibition of glutamatergic mdPons neurons decreased both the sniffing and pDA responses to novel odors (n=43 cells). Red frame: in remaining 4 animals, the same manipulation increased both sniffing and pDA activity (n=64 cells). B) Comparison of viral expression between the animals in which optogenetic inhibtion decreased (blue) vs. increased (red) orienting. The animals in which inhibition increased sniffing had more expression in the poserio-medial area (see AP – 4.45 & -4.55mm). The additional expression involved mainly 3 regions: central gray (alpha and beta parts), sphenoid nucleus and trochlear nucleus. Because stimulation with red light consistently increased both sniffing and pDA activity in both subgroups, these results were pulled together and illustrated on Fig. 5.

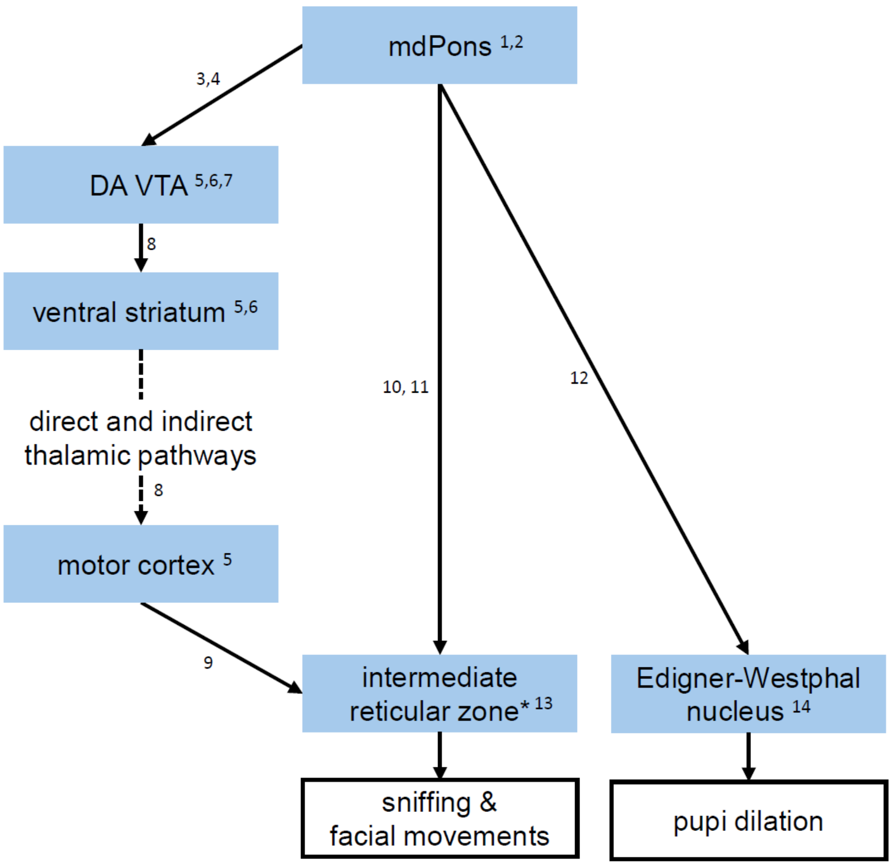


**Supplementary Figure 9.** Proposed anatomical circuit supporting active sensory exploration. Numbers next to brain regions indicate evidence that stimulating a given area evokes sniffing and/or facial movements and/or pupil dilation. Numbers next to arrows indicate evidence for the existence of anatomical connections between regions.

*Intermediate reticular zone is a collection of densely interconnected nuclei that control breathing, sniffing, facial movements etc. in a synchronized manner. It contains such areas as preBotzinger nucleus (which controls respiratory rate) and facial nucleus (which controls facial movements). [12]
